# Next-generation ABACUS biosensors reveal cellular ABA dynamics driving root growth at low aerial humidity

**DOI:** 10.1101/2022.10.19.512731

**Authors:** James Rowe, Mathieu Grangé-Guermente, Marino Exposito-Rodriguez, Rinukshi Wimalasekera, Martin Lenz, Kartika Shetty, Sean R. Cutler, Alexander M. Jones

## Abstract

The plant hormone abscisic acid (ABA) accumulates under abiotic stress to recast water relations and development. To overcome a lack of high-resolution, sensitive reporters, we developed ABACUS2s, next-generation FRET biosensors for ABA with high affinity, signal-to-noise ratio and orthogonality, that reveal endogenous ABA patterns in Arabidopsis thaliana. We mapped stress-induced ABA dynamics in high-resolution to reveal the cellular basis for local and systemic ABA functions. At reduced foliar humidity, roots cells accumulated ABA in the elongation zone, the site of phloem transported ABA unloading. Phloem ABA and root ABA signalling were both essential to maintain root growth at low humidity. ABA coordinates a robust system to maintain root growth in response to foliar stresses, enabling plants to maintain foraging of deeper soil for water uptake

## Main Text

Plant decision making is distributed rather than centrally coordinated, but to survive and overcome stresses such as a lack of water, responses must also be systemically coordinated. Abscisic acid (ABA) is a phytohormone that accumulates systemically under various local water stresses to coordinate responses over a complex and often-large morphology. When roots experience low-water stress, for example, ABA closes the microscopic pores on leaves (stomata), to limit systemic water loss ^1–3^. Interestingly, leaf water loss can cause changes in root growth responses and architecture - increasing transpiration genetically or through increased airflow produces larger root systems in *Arabidopsis* ^4^ and low relative humidity (RH) can promote root growth in many species ^5–7^. Although a molecular mechanism remains elusive, it has been proposed that ABA, acting as a dehydration signal, could be coordinating these root growth responses ^4,8^. The sites of ABA biosynthesis, metabolism and translocation are the subject of intensive research, but progress has been hampered by limitations in tools to quantify accumulation and depletion of ABA on a tissue/cellular scale where regulatory decisions controlling ABA dynamics are made ^9,10^. The availability of sensitive reporters, particularly Förster Resonance Energy Transfer (FRET) biosensors, for hormones, second messengers and metabolism are revolutionizing plant development, signalling and photosynthesis research ^11^. Such biosensors are powerful tools to quantify metabolites *in vivo* at high spatiotemporal resolution ^11^, including phytohormones under changing environmental conditions ^12–16^. Direct ABA FRET biosensors that do not require additional signalling components have broad application potential beyond ABA quantification in plant cells and sub-cellular compartments. For example in ABA synthesizing pathogenic fungi ^17^, in human granulocytes where ABA is a cytokine^18^, or in extracts from organisms where genetic modification is difficult using purified protein *in vitro*^19^. However, existing ABA FRET biosensors, ABAleons and Abscisic Acid Concentration and Uptake Sensors 1 (ABACUS1s) ^13,14,20^, lack the full complement of strengths required to easily quantify ABA. Therefore, we engineered next-generation ABA biosensors and deployed them to dissect cellular ABA dynamics and mobilization in response to foliar humidity stress and to establish a systemic role for ABA to maintain local root growth in response to a distant shoot stress.

In ABAleons and ABACUS1 biosensors, ABA sensory domains are connected by linkers to a pair of fluorescent proteins (FP) (Extended Data Fig.1). The orientation and distance between these FPs determines the transfer of excitation energy via FRET from a donor FP to an acceptor FP. Ligand-induced conformational changes in sensory domains alter the relative positions of the FPs, which can be detected by exciting the donor and measuring a change in relative acceptor and donor emissions, hereafter referred to as emission ratio change.

ABAleons are sensitive to endogenous ABA concentrations, but have poor signal-to-noise ratios (small emission ratio change). ABACUS1s have high signal-to-noise ratio but poor sensitivity for endogenous ABA ^13,14^. Ideal biosensors are also orthogonal with minimal interaction with endogenous signalling. ABAleons have strong ABA hyposensitivity phenotypes while ABACUS1s have minor ABA hypersensitivity phenotypes ^13,14,21^. We used ABACUS1-2μ as the basis to engineer next-generation biosensors with high sensitivity, emission ratio change and orthogonality (Extended Data Fig.2).

ABACUS1-2μ has a K_D_(ABA) of ∼2 μM and consists of an N-terminal FRET acceptor (edCitrine), an attB1 linker, a sensory domain consisting of a mutated **PY**RABACTIN RESISTANT 1 **L**IKE **1** (**PYL1 H87P**) ABA receptor and a truncated PROTEIN PHOSPHATASE 2C (PP2C) co-receptor, **AB**SCISIC ACID **I**NSENSITIVE 1 **a**ba **i**nteracting **d**omain (**ABI1aid**), an attB2 linker, and a C-terminal FRET donor (edCerulean) ^14^. We introduced a second PYL1 mutation (A190V) into ABACUS1-2μ that is known to increase ABA affinity of PYL1 ^22^. The resulting ABACUS had increased affinity but reduced emission ratio change *in vitro* (Fig 1. a, b, Extended Data Fig.2, Extended Data Table 1).

**Fig. 1.**
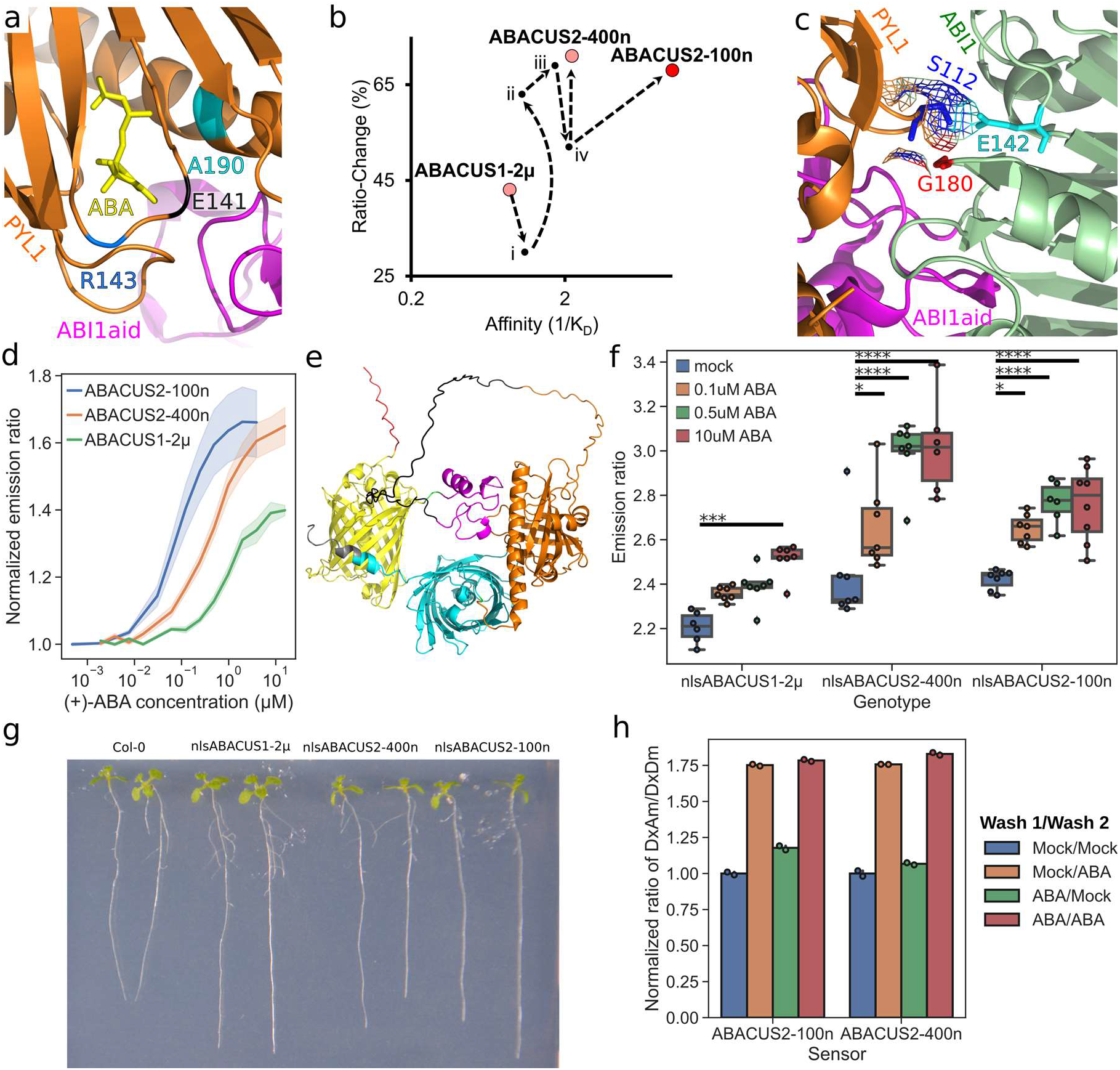
ABACUS2-100n and ABACUS2-400n offer higher ratio change and affinity than ABACUS1 and moderate phenotypes. a) Location of binding site mutations ABA binding site (A190V, E141D, R143S) mapped onto crystal structure PDB: 3JRQ. b) Affinity vs ratio change of ABACUS variants. Intermediate sensor versions are as follows: i. ABACUS1-2μA190V, ii. ABACUS1-2μA190V PPP-L52-P linkers iii ABACUS1-2μA190V PPP-L52-P linkers S112A, iv ABACUS1-2μA190V, PPP-L52-P linkers, S112A, edCitrineT9 edCeruleanT7 c) The ABI1-PYL1 interface. S112 (blue) interacts with residues in ABI1 (light green) but not the ABIaid (magenta). From crystal structure PDB: 3JRQ. d) Purified protein emission ratio titration of (+) - ABA for ABACUS variants. Line indicates mean of multiple independent extractions and titrations, shaded region indicates the standard error of the mean. ABACUS1-2μ (n=6), ABACUS2-400n (n=16) and ABACUS2-100n (n=13). e) Illustrative Collabfold/Alphafold MMseqs2prediction of ABACUS2-100n structure. Domains are: nuclear localization signal (red), edCitrineT9 (yellow) ABI1aid(ABI1 49 aa truncation, magenta), PYL1(H87P, S112A, A190V, E141D), L52 linker (black), T7edCerulean (cyan) myc tag (dark red). Structural alignment with PDB: 3JRQ of ABA-PYL1-ABI1 is available in the supplement. f) nlsABACUS emission ratio responses in *Arabidopsis* roots exposed for 30 minutes to various concentrations of ABA. Each point indicates median nuclear emission ratio for an individual root z-stack. Representative images are available in Extended Data Fig. 3. 2-way ANOVA, (sensor: F=64.9, P<0.0001, DF=2; Treatment F=37.91, P<0.0001, DF=3; Interaction: F=3.349, P=0.0059 DF=6) Asterisks indicate significance with a Tukey post hoc test *:p<0.05, **:p<0.01, ***:p<0.001, ****:p<0.0001, n=6,7,8,7,8,7,6,6,7,7,6,8 respectively g) Visual phenotypes at 11 days after stratification (DAS) of nlsABACUS1-μ, nlsABACUS2-400n line 7 and nlsABACUS2-100n line 7 h) *in vitro* reversibility testing of purified ABACUS2-400n and ABACUS2-100n sensors. n=2

Engineering increased emission ratio change is semi-empirical as mutations in any moiety may boost the transduction of ligand-binding into FRET change, but a first target is often the linkers between sensory domain and the FRET pair ^23^. Replacing the attB linkers with shorter, less flexible, proline linkers rescued emission ratio change of the A190V mutant (Fig 1b, Extended Data Fig.2, Extended Data Table 1).

A higher-affinity PYL1 receptor would likely exacerbate ABA hypersensitivity phenotypes of ABACUS expressing plants ^14^. Therefore, we introduced an orthogonalizing mutation, PYL1 S112A (Fig 1b, c), to reduce PYL1 signalling through endogenous co-receptors (e.g. ABI1, ABI2, and HAB1) ^24^. Because this mutation disrupts PYL1 interaction with ABI1 residues E142 and G180 that are absent in the ABI1aid truncation of ABACUSs (Fig 1c), we correctly predicted PYL1 S112A would not lower emission ratio change or affinity (Fig 1b, Extended Data Fig.2).

We next incorrectly predicted that truncating the flexible fluorescent protein termini facing the sensory domain (edCitrineT9, T7edCerulean) would increase ratio change further (Fig 1b, Extended Data Fig.2, Extended Data Table 1). Nonetheless, emission ratio change could be restored along with further affinity improvements by introducing either of two separate mutations to a PYL1 region – the “latch” - that is important for both PYL1-ABA and PYL1-PP2C interactions ^25^. We selected the first mutation, PYL1 R143S, to alter water mediated PYL1-ABA-ABI1aid interactions. This produced our highest ratio change biosensor that has ABA sensitivity suitable for *in planta* studies, which we named **ABACUS2-400n** (K_D_ (ABA): 445 nM, *in vitro* emission ratio change: +71%, Fig 1 b, d, Extended Data Fig. 1, 2, Extended Data Table 1). The second mutation, PYL1 E141D, inspired by sequences of the high-affinity PYL8 and PYL9 ABA receptors, produced a high ratio-change sensor with our highest affinity, which we named **ABACUS2-100n** (K_D_(ABA): 98 nM, *in vitro* emission ratio change: +67%, Fig 1b, d, e, Extended Data Fig 1, 2, 3, Extended Data Fig 1). *In vitro* assays against other phytohormones, salts and ABA related compounds demonstrated that ABACUS2 sensors are highly for specific for ABA and the ABA agonist Pyrabactin (Extended Data Fig 4).

Improved promoter/terminator combinations allowed us to express ABACUS2 sensors with nuclear localization signals (nls) in wildtype *Arabidopsis thaliana* (Col-0), overcoming our previously severe ABACUS1 silencing problems ^14^. Nuclear localization allows easy discrimination of the fluorescence of neighbouring cells and the exclusion of non-nuclear background and auto-fluorescence during image processing ^11^. To accelerate this image processing, we developed a comprehensive image analysis toolset to quickly analyse confocal stacks in 3D/4D, allowing us to robustly quantify and visualize nuclear emission ratios within moments (See supplemental methods and ^26^).

In Col-0, nlsABACUS2-400n and nlsABACUS2-100n respond strongly at lower concentrations of exogenous ABA than ABACUS1-2μ (Fig 1f, Extended Data Fig. 5), confirming their improved sensitivity *in planta*. The ABACUS2 emission ratio changes are significantly larger than ABACUS1-2μ^14^ or other state-of-the-art ABA sensors (ABAleonSD1-3L21) ^21^ (Fig 1f, Extended Data Fig. 5, 6). Even though our new nlsABACUS2 lines had 5 – 25-fold higher affinity than previous nlsABACUS1-2μ lines, phenotypes are relatively mild (Fig 1g, Extended Data Fig. 7, 8). Without exogenous ABA, nlsABACUS-400n germinates normally, and nlsABACUS2-100n is slightly delayed (Fig. 1g, Extended Data Fig. 7), however both display robust post germination root growth. With ABA, nlsABACUS2 lines display a hypersensitive germination inhibition (Extended Data Fig. 7), but wildtype-like root growth (Extended Data Fig. 8) suggesting that the ABACUS2 PYL1 is likely somewhat active *in planta*, but the PYL1 S112A orthogonalizing mutation successfully reduced ABACUS2 PYL1 interaction with endogenous PP2Cs.

Both ABACUS2 sensors were rapidly reversible *in vitro* (Fig 1 h) and nlsABACUS2-400n emission ratios decreased rapidly following a 50 μM ABA pulse (Fig 2a, b) delivered to roots growing in the RootChip microfluidics system ^12,27^.

**Figure 2.**
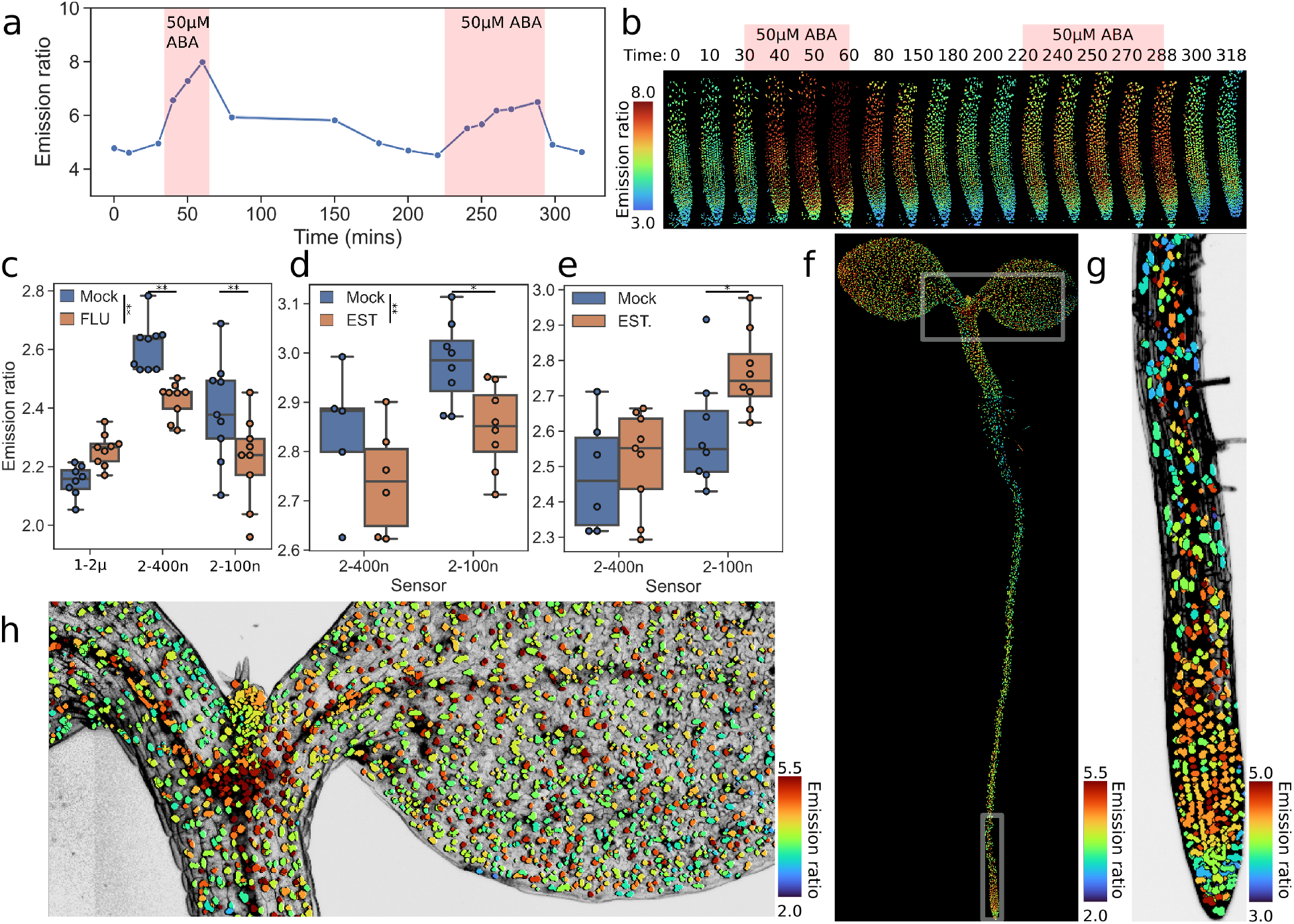
ABACUS2-100n and ABACUS2-400n reveal endogenous ABA patterns, accumulations and depletions. a) Graph and maximum intensity Z projection (b) of emission ratios of ABACUS2-400n roots responding to 50μM exogenous ABA treatment pulses, performed with the RootChip microfluidics system. Number of nuclei in each time point respectively: 996,1036,996, 856, 1020, 931, 875, 832, 935, 931, 974, 924, 931, 1003, 932, 972, 963, 1002 c) 24h fluridone treatment effect on emission ratios of nlsABACUS roots. Representative images in Extended Data Fig. 9. 2-way ANOVA Treatment: F=7.4, p=0.009 DF=1, Sensor F=38.0 p<0.0001, DF=2, Interaction F=9.7 p=0.0003, DF=2. n=8,9,9,9,9,9 respectively A Tukey post hoc test was used for multiple comparisons. d) 24 hour catabolism induction (10μM Estradiol, UBQ10::XVE:CYP707A3) reduced nlsABACUS2-100n and nlsABACUS2-400n emission ratios in Arabidopsis roots. Representative images in Extended Data Fig. 10. 2-way ANOVA Treatment: F=8.1, p=0.009 DF=1, Sensor F=9.9 p<0.0046 DF=1, Interaction F=0.2 p=0.660. DF=1 n= 5,6,8,8 respectively A Tukey post hoc test was used for multiple comparisons e) 24 hour biosynthesis induction (5μM Estradiol, UBQ10::XVE:NCED3) increased nlsABACUS2-100n emission ratios in Arabidopsis roots. Representative images in Extended Data Fig. 11. Each point indicates mean nuclear emission ratio for an individual root z-stack. 2-way ANOVA Treatment: F=4.0, p=0.055, DF=1, Sensor F=4.84 p=0.037, DF=1, Interaction F=2.0 p=0.167, DF=1. n=6,9,8,8 respectively. A Tukey post hoc test was used for multiple comparisons. f) Nearest point Z-projection of whole plant nlsABACUS2-400n emission ratios. Boxes indicate approximate crops of g and h. g) Nearest point Z-projection of nlsABACUS2-400n ratios in the root tip. Gray to black channel represents propidium iodide counterstaining of cell walls. h) Nearest point Z-projection of ABACUS2-400n ratios in the cotyledons and hypocotyl. Gray to black channel represents propidium iodide counterstaining of cell walls. Asterisks indicate statistical significance *:p<0.05, **:p<0.01, ***:p<0.001, ****:p<0.0001

To determine if the increased affinity of ABACUS2s allows them to reliably measure endogenous variations in ABA levels, unlike the lower sensitivity ABACUS1 sensors, we undertook a pharmacological and inducible-genetics approach. The ABA biosynthesis inhibitor fluridone reduced nlsABACUS2 emission ratios, but nlsABACUS1-2μ remained level (Fig 2c, S9). Inducing ABA catabolism with CYTOCHROME P450, FAMILY 707, SUBFAMILY A, POLYPEPTIDE 3 (CYP707A3 ^28^) overexpression reduced nlsABACUS2 emission ratios (Fig 2d, Extended Data Fig. 10), and inducing ABA biosynthesis with 9-CIS-EPOXYCAROTENOID DIOXYGENASE 3 (NCED3) overexpression increased emission ratios (Fig 2e, Extended Data Fig. 11). Therefore, nlsABACUS2 sensors respond to physiological levels of ABA.

The availability of sensitive reporters for other phytohormones such as auxin revolutionized plant developmental biology, by revealing localized activity of a key hormone for morphogenesis and patterning ^11^. Similarly, sites of ABA accumulation may give insights into developmental regulation and stress responses. Therefore, we used nlsABACUS2s to determine the distribution of ABA in *Arabidopsis* plants (Fig 2f, g, h, Extended Data Fig. 12). nlsABACUS2 seedlings had higher emission ratios in internal tissues of the cotyledons and hypocotyl, including the vasculature, indicating high ABA in these tissues.

High ABA in the shoot vasculature is significant, as the phloem companion cells are a key site for ABA biosynthesis ^29^ and ABA is thought to be transported in the phloem ^8^. The phloem transports sugars, hormones and other metabolites from shoot to root, where it can be unloaded via the phloem-pole pericycle cells in the root elongation zone from two distinct vascular poles ^30^. nlsABACUS2 roots show high emission ratios in these tissues, (Fig 2g, Extended Data Fig. 12) so we used Single Plane Illumination Microscopy (SPIM) to examine whether phloem sourced ABA is unloaded here (Fig 3a, b). Before treatment, nlsABACUS2-400n emission ratios were higher in two poles of the root vasculature, as would be predicted for a phloem-transported hormone (Fig 3b.). Root emission ratios increased rapidly following shoot ABA treatment, starting in vascular poles, then spreading radially through the elongation zone and longitudinally to the differentiation zone and mature root (Fig 3a, b) – matching patterns of when shoot applied fluorescent dyes are unloaded from the phloem ^30^.

**Fig 3.**
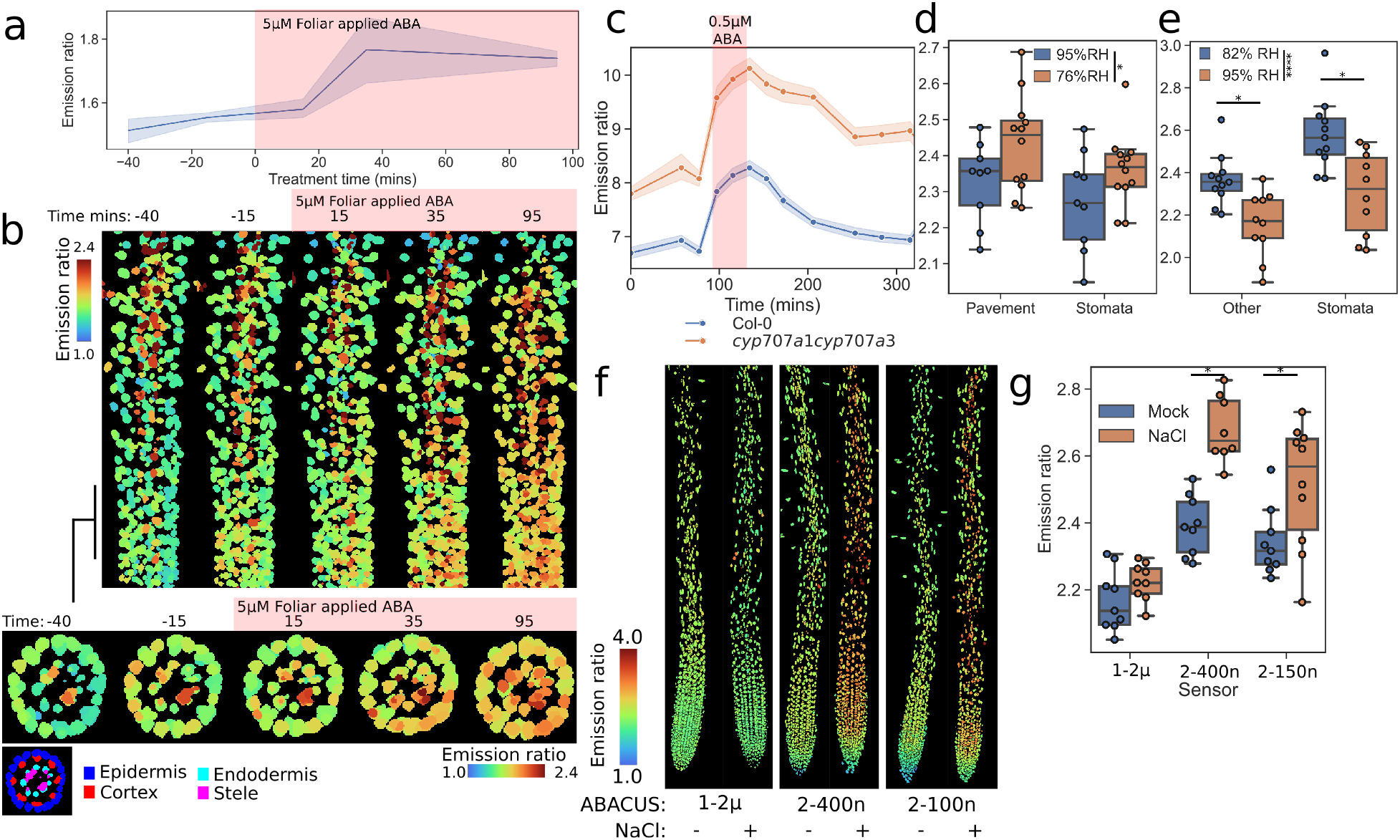
ABA levels are high in the internal tissues of the cotyledons, the vasculature, and the root elongation zone. Leaves and roots respond to local abiotic stresses with ABA accumulation. a) SPIM microscopy of nlsABACUS2-400n exposed to a 5μM ABA treatment to the foliar tissues. Roots are isolated from the foliar tissues so emission ratio increases must come from ABA transport. Number of nuclei in each time point respectively: 458,488,476,481,466 b) Max Z projection and a Max Y projection of the indicated area of the data quantified in a). c) nlsABACUS2-400n in Col-0 and *cyp707a1cyp707a3* emission ratios under ABA pulsing. N=9, 12, 9, 12 respectively d) nlsABACUS2-400n emission ratios increase in response to a 6 hour humidity decrease. Relative humidity (RH) indicates the measured humidity at leaf height during the treatments. Representative images and peristomatal distance are available in Extended Data Fig. 11. 2-way ANOVA Humidity F=6.29 p=0.0165 DF=1, Cell type F=2.08 p=0.157 DF=1, Interaction F=0.0088 p=0.926 DF=1. n=11, 10, 11, 10 respectively. A Tukey post hoc test was used for multiple comparisons e) nlsABACUS2-400n emission ratios decrease in response to a 6 hour humidity increase. RH indicates the measured relative humidity at leaf height during the treatments. Representative images and peristomatal distance are available in Extended Data Fig. 12. Each point indicates median nuclear emission ratio for an individual z-stack. 2-way ANOVA (Treatment F=24.1, p<0.0001 DF=1; Cell type F=13.14, p= 0.0008 DF=1; interaction F=0.46 p=0.498 DF=1). A Tukey post hoc test was used for multiple comparisons f) and ABACUS2 emission ratios in response to 6 hours 100mM NaCl treatment. 2-way ANOVA (Treatment F=30.6, p<0.0001 DF=1; Cell type F=41.02, p< 0.0001 DF=2; interaction F=4.43 p=0.017 DF=2). n=9, 9, 9, 8, 9, 10. A Tukey post hoc test was used for multiple comparisons. Asterisks indicate statistical significance *:p<0.05, **:p<0.01, ***:p<0.001, ****:p<0.0001

Exogenous ABA causes concentration-dependent promotion or inhibition of root growth ^31^, so ABA from the phloem must be tightly regulated independently of local biosynthesis. The abscisic acid 8’-hydroxylases CYP707A1-4 catabolic enzymes have been implicated in eliminating ABA after stress ^32,33^. *CYP707A1* and *CYP707A3* are the isoforms most expressed in the root ^28^ and *cyp707a1cyp707a3* double mutants ^34^ displayed a strong over-accumulation of ABA in the root tip (Extended Data Fig. 13). Exogenous ABA pulsing revealed larger emission ratio increases in *cyp707a1cyp707a3*, and considerably slower elimination than Col-0 (Fig 3c). Whilst these enzymes are critical to prevent over-accumulation of ABA in the root tip, other ABA depletion mechanisms must also contribute to the ABA elimination as there is still a slow reduction in *cyp707a1cyp707a3* nlsABACUS2-400n emission ratios following an ABA pulse (Fig 3c).

ABA has numerous roles protecting plants from abiotic stress, particularly osmotic and ionic stresses. During salt stress, root ABA responses mediate endodermal cell wall suberization ^35,36^, limiting ion and water flow to protect the plant, however it’s currently unclear which cells accumulate ABA. High-resolution imaging of ABACUS2-400n gave us an unparalleled view of the ABA accumulation after a six-hour 100mM NaCl stress (Fig 3f, g, Extended Data Fig. 14) allowing us to quantify which tissues accumulate ABA. Under salt stress, the stele (a site of ABA biosynthesis) and endodermis (a site of ABA dependent protective responses) of the differentiation/maturation zones accumulated more ABA than the surrounding epidermis and cortex tissues (Fig Extended Data Fig. 14).

Confident that we could image and detect cell type specific ABA accumulations, we decided to investigate the effect of humidity on plant ABA levels and responses in detail. A six-hour humidity drop increased emission ratios in stomata and pavement cells expressing nlsABACUS2-400n (Fig 3d, Extended Data Fig. 15), which coincided with a decreased stomatal aperture (Extended Data Fig. 15). Leaf humidity increases trigger expression of ABA catabolic genes *CYP707A1* and *CYP707A3* ^33^ and nlsABACUS2-400n emission ratios decreased following a humidity increase, and stomata opened (Fig 3e, Extended Data Fig. 16). Remarkably, nlsABACUS2-400n emission ratios responded similarly in pavement cells and stomatal cells to humidity changes (Fig 3e, Extended Data Fig. 16). ABA famously closes stomata, and along with the vasculature, stomata have been proposed as sites of ABA biosynthesis ^29,33,37^, but little attention has been paid to whether pavement cells accumulate ABA. Such broad ABA increases may indicate a systemic response that travels beyond the tissues responsible for fast local responses.

As foliar ABA levels increase following a humidity stress and foliar ABA can be transported to the root (Fig 3a,b) ^8,38^, we predicted that a local shoot stress may cause ABA accumulation in roots, affecting root growth and development. Leaf transpiration rates can affect root growth and morphology though an uncharacterized mechanism ^4^, however root plasticity is strongly ABA regulated under salt and other local water stresses ^39,40^.

We developed a system where leaves could be exposed to low humidity and roots would remain hydrated (Fig Extended Data Fig. 17) and maintain robust root growth (Fig 4a). Remarkably, the ABA biosynthesis mutant *aba2* suffered a strong root growth inhibition under low humidity (Fig 4A) implying that ABA signalling functions to maintain root growth when foliar humidity is low, a scenario common in irrigation agriculture.

**Fig 4.**
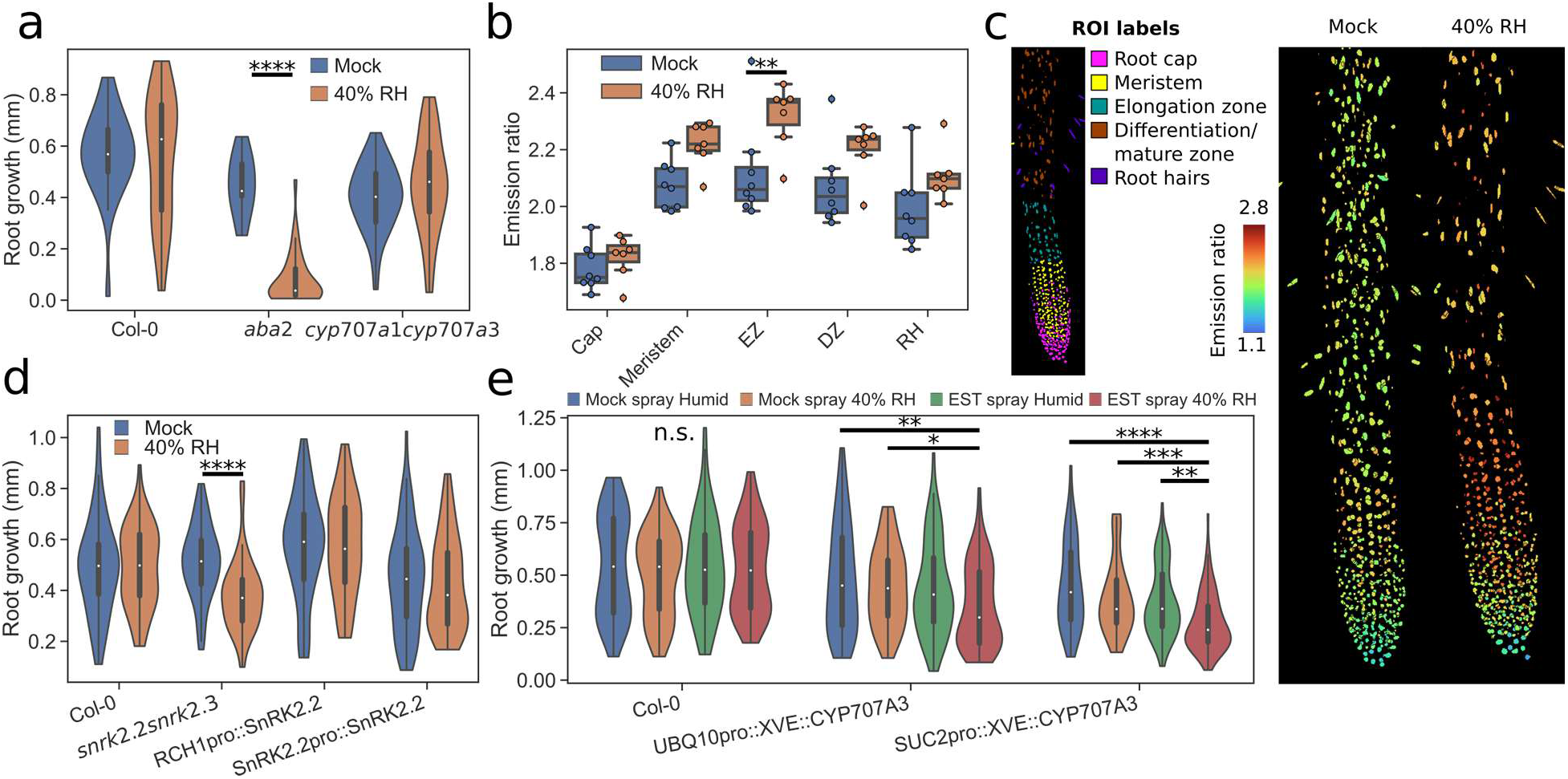
Foliar humidity decreases induce root ABA accumulation to regulate root growth. a) Root growth of 6 DAS Col-0, *aba2* and *cyp707a1cyp707a3* in response to 7 hour foliar humidity drop. 2-way ANOVA (Treatment F=15.0, p=0.002 DF=1; Genotype F=31.8, p< 0.0001 DF=2; interaction F=16.7 p<0.0001 DF=2). n=31,27,36,9,35,40 respectively. A Tukey post hoc test was used for multiple comparisons. b) and c) Root emission ratios of nlsABACUS2-100n increase under humidity stress, with the elongation zone showing a significant ABA accumulation and little response in the root cap. EZ: Elongation zone, DZ: Differentiation/maturation zone, RH: Root hair. Each point indicates median nuclear emission ratio for an individual root z-stack. 2-way ANOVA (Treatment F=23.64 DF=1 p<0.0001; Root zone F=31.29 DF=4, p< 0.0001; interaction F=0.978 DF=4 p=0.426).n=8,7,8,7,8,7,8,7,8,7 respectively. A Tukey post hoc test was used for multiple comparisons. d) Root growth of 6 DAS Col-0, *snrk2*.*2snrk2*.*3, snrk2*.*2snrk2*.*3* RCH1pro::SnRK2.2, and *snrk2*.*2snrk2*.*3* SnRK2pro::SnRK2.2 in response to a short term foliar humidity drop. 2-way ANOVA (Treatment F=5.158 p=0.0235 DF=1; Genotype F=16.68, p< 0.0001 DF=3; interaction F=4.303 p=0.00052 DF=3). n= 58,64,80,60,69,64,77,52 respectively. A Tukey post hoc test was used for multiple comparisons. e) Root growth of 6 DAS Col-0, UBQ10pro::XVE::CYP707A3 and SUC2pro::XVE::CYP707A3 in response to a short term foliar humidity drop 24 hours after a shoot spray of 50 μM β-Estradiol. 2-way ANOVA (Treatment F=10.91 p<0.0001 DF=3; Genotype F=51.57, p< 0.0001 DF=2; interaction F=3.063 p=0.0057 DF=6). n=81,90,70,70,58,70,67,68,116,67,85,111 respectively A Tukey post hoc test was used for multiple comparisons. Asterisks indicate statistical significance *:p<0.05, **:p<0.01, ***:p<0.001, ****:p<0.0001

nlsABACUS2-100n roots displayed increased root emission ratios at low humidity, which were particularly prevalent in the elongation zone, the site of phloem unloading and a tissue critical for root growth (Fig 4b, c). We took a targeted genetic approach to determine if increases in root ABA are critical for plants to increase/maintain root growth at low humidity. ABA responses rely on the activity of the SnRK2 kinases SnRK2.2, SnRK2.3 and SnRK2.6/OPEN STOMATA 1(OST1) which phosphorylate downstream transcription factors to activate gene expression ^41^. *snrk2*.*2snrk2*.*3* mutants have ABA insensitive roots but can maintain normal stomatal function and closure responses under stress due to a functional SnRK2.6 protein, the principal SnRK2 responsible for phosphorylating ion channels to close stomata ^41,42^.

Like *aba2*, the *snrk2*.*2snrk2*.*3* mutant demonstrated a reduced root elongation rate under humidity stress (Fig 4d). Complementation of the *snrk2*.*2snrk2*.*3* mutant specifically in the root tip with RCH1pro::SnRK2.2 ^43^ allowed plants to maintain root growth under a humidity stress, indicating that local ABA signalling is required to regulate root growth as humidity varies (Fig 4d).

ABA synthesized in the phloem companion cells ^29^, is likely to be transported to phloem sinks including the root elongation zone ^30^. We posited that the root induction of ABA accumulation at low foliar humidity might be phloem sourced so performed targeted ABA depletions by controlled induction of *CYP707A3* overexpression (Fig 4e). Whether ectopic ABA depletion was restricted to phloem-loading companion cells (SUC2pro::XVE::CYP707A3) or ubiquitous (UBQ10pro::XVE::CYP707A3), root growth was inhibited at low foliar humidity (Fig 4e). Taken together, our results indicate that phloem ABA and root tip ABA signalling regulate root growth during a distal humidity stress in leaves.

A series of local and systemic responses are required for plants to respond to varying water availability. Phenotypic data suggests that plant roots can respond to local osmotic differences through ABA, for example growing towards water (hydrotropism) ^43^, but determining if ABA levels vary across a root has been experimentally challenging. nlsABACUS2 allows local increases in ABA to be visualized at the cellular level, as in the accompanying submission. Mehra *et al*. show a local increase in root ABA in response to root growth through air spaces, without an increase in foliar ABA levels^44^. Similarly, we have shown that salt stress induces ABA accumulation in the tissues where a protective response is required, the root endodermis. However, plant roots can also induce systemic ABA accumulation. During soil drying, both sulfate and CLE25 peptides can be transported from the root to induce foliar ABA accumulation, closing stomata and limiting water loss ^1–3^. During drought, some of this shoot derived ABA is also transported down to the root to promote and maintain root growth, allowing more access to soil water ^8^. That foliar tissues can sense water loss has long been known, as plants quickly regulate their stomatal aperture in response to an increased vapor pressure deficit, a process enhanced by foliar ABA accumulation ^45^. Here we show, with cellular resolution afforded by nlsABACUS2 biosensors, that foliar drying can also regulate root ABA accumulation and that this root ABA is important to maintain root growth under stress. This demonstrates that the root and shoot can each systemically regulate each other’s responses to stresses that may only be experienced locally, providing a robust system to maintain plant water status.

## Methods

### Data visualization and statistical analysis

Unless otherwise stated, data was processed as Pandas dataframes in Python, using statsmodel/Graphpad prism for statistics and Seaborn/matplotlib/excel for plotting.

### Generation of ABACUS affinity and orthogonality variants

Single amino acid mutations of the PYL1 domain of ABACUS1 in the pDRFLIP38-ABACUS1-2μ vector ^14^ using the QuikChange II XL [Agilent] site-directed mutagenesis kit according to manufacturer’s instructions. All primers used for site-directed mutagenesis are listed in **Extended Data Table 2**.

### Generation of ABACUS ratio-change variants

The edCitrine present in ABACUS1 variants ^14^ was exchanged with a codon diversified version for optimal expression in yeast and to allow PCR based cloning methods. The synthetic DNA fragment containing the codon diversified edCitrine was introduced in the ABACUS yeast expression vectors using the In-Fusion kit [Takara Bio] according to manufacturer’s instructions.

The poly-proline screen variants, which included substitution of the attB1 and attB2 linkers of ABACUS1 with 1-3 proline residues, and the fluorescent proteins truncations were obtained using the In-Fusion kit according to manufacturer’s instructions. All primers used for In-Fusion cloning are listed in **Extended Data Table 2**.

### Fluorescence analysis and titration with (+)-ABA of protein purified cell lysate

Yeast cell cultures (OD600 ≈0.6) containing yeast expression vector pDRFLIP38-ABACUS1-2μ or variants were centrifuged at 4000g for 10min, washed once in 1 mL of 50 mM MOPS Buffer (pH7.4), transferred to 1.5 mL micro-centrifuge tubes and centrifuged again at 10000g for 1min. The supernatant was discarded and 1 mL of chilled glass bead slurry (50 mM MOPS pH7.4, 0.1% Triton X-100 and 50% vol/vol 0.5 mm Zirconia/Silica beads [Thistle Scientific]) was added to the yeast pellet inside each tube. The tubes were then vortexed at maximum power at 4°C for 5min. The tubes were then centrifuged at 14000x at 4°C for 10 min. The supernatant was transferred to previously prepared HisPur Cobalt Spin Columns, 0.2 mL [Thermo Fisher Scientific]. Protein purification was performed following manufacturer’s instructions. The subsequent first elution from the purification column was diluted in 50mM MOPS solution. The tubes were briefly vortexed and 100μl of diluted eluate was transferred to 96-well flat bottom clear microplate [Greiner]. A serial dilution of (+)-ABA [Cayman Chemical] was made using a 4.5mM stock solution in ethanol and sequentially diluting it in 50mM MOPS solution. 50μL of each (+)- ABA dilution was added to 100μL of sensor eluate. The sample’s fluorescence emission was recorded using a SpectraMax i3x [Molecular Devices], scanning from 470 to 550nm after excitation at 430nm with a bandwidth of 5nm. The data produced was analysed using GraphPad Prism [GraphPad Software] to determine the k_D_ and ratio change of each sensor, assuming the Hill function with a single binding site.

### Structure prediction

nlsABACUS2-100n structures were predicted (for illustrative purposes only) using the Colab-Fold 1.4 notebook, based on Alphafold2, using MMseqs2 for homology detection and multiple sequence alignment pairing ^46^. The highest ranked (by pLDDT) prediction was used. Structural validation and confidence measures are shown in Extended Data Fig. 3.

### Cloning ABA biosynthetic and catabolic enzyme constructs for inducible expression in plants

*AtNCED3* (AT3G14440.1) was amplified with attB1/attB2 sites with q5 polymerase, following manufacturer’s instructions, and inserted into pDONR221-f1 ^47^ with a BP reaction. *AtCYP707A3* (AT5G45340.1) coding sequence with attL1/attR1 sites was synthesized in pUC19 from Genewiz. These could then be combined with *p1R4-pAtSUC2:XVE/p1R4-pUBQ10:XVE* and *p2R3a-NosT* ^48^ through a Multisite LR reaction to generate *SUC2pro::XVE>>CYP707A3, UBQ10pro::XVE>>CYP707A3*, and *UBQ10pro::XVE>>NCED3* in *pHm43GW* ^49^. Gateway cloning was performed following manufacturer’s instructions.

### Cloning ABACUS2 constructs for expression in plants

*ABACUS2-100n* and *ABACUS2-400n* were subcloned from the yeast vectors, reverting the codon diversification of the edCitrineT9. To do this, the sensory domain to the stop codon were amplified with attB1/attB2 sites and inserted into *pDONR221-f1* with a BP reaction. *nls-edCitrineT9* was amplified from *nlsABACUS1-2μ* plasmid ^14^ and introduced into the *pENTR221-f1-ABACUS2-truncation* vectors using In-Fusion cloning (Takara) to generate *pENTR-nlsABACUS2-100n* and *pENTR-nlsABACUS2-400n*. ABACUS2 Gateway entry clones were combined with *p1R4-pUBQ10* and *p2R3a-NosT* into *pFR7m34GW* ^50^ through a multisite LR reaction. Primers are listed in **Extended Data Table 2**.

### Plant transformation

*Arabidopsis thaliana* plants (Columbia, Col-0 background) were transformed by the floral dip method ^51^ and successful transformants were identified by FAST RED screening ^52^, or hygromycin selection. Full details of *Arabidopsis* germplasm are available in Extended Data Table 3

### Plant growth conditions

For endpoint root imaging experiments, plants were grown under long day conditions (110 μE, 22 °C 18hrs, 0 μE, 18°C 6hrs).

### Salt treatment

Seeds were surface sterilized with 96% ethanol, then sown on ½ Murashige and Skoog (MS) ^53^ 0.05% MES plates pH 5.7, sealed with micropore tape, then stratified for 4 days at 4°C. Plants were grown for 5 DAG before a 5.5 hour treatment. Treatment consisted of a transfer to ½ MS plates containing 100mM (45511 Merck) or a fresh ½ MS MES plate for mock.

### Fluridone treatment

Seeds were surface sterilized with 96% ethanol, then sown on ½ MS plates 0.05% MES pH 5.7, sealed with micropore tape, then stratified for 4 days at 4°C. Plants were grown for 5 DAG before a 24 hour treatment. For treatment, plants were transferred to ½ MS plates containing 0.4 μM fluridone (45511 Merck), or an Ethanol mock.

### β- Estradiol induction of ABA biosynthesis/catabolism

Seeds were surface sterilized with 96% ethanol, then sown on ½ MS 0.05% MES plates pH 5.7, sealed with micropore tape, then stratified for 4 days at 4°C. Plants were grown for 5 DAG before a 24 hour treatment. Treatment consisted of a transfer to ½ MS 0.05% MES plates pH 5.7 containing 10μM β-Estradiol or a DMSO mock.

### Leaf humidity treatments for leaf imaging

nlsABACUS-400n seeds were surface sterilized with 96% ethanol, then stratified for 4 days at 4°C in sterile deionized water, before sowing on F2 Levington’s compost. Plants were grown (120 μE, 22 °C 18hrs, 0 μE, 18°C 6hrs) for 15 DAG before humidity treatment. Plants were germinated under a clear plastic propagator lid, which was removed at 4 DAG.

For a humidity increase, the chamber was set to 60% RH, and humidity increased by placing a propagator lid over the plants for 6 hours before imaging. Humidity and temperature were measured at leaf height above compost at ∼95% RH 22 °C for treatment, and ∼82% RH 22 °C for mock. Humidity and temperature were measured using a BME280 sensor.

For a humidity decrease, the chamber was set to 40% RH, and were grown with a propagator lid until treatment. For treatment, compost was covered with acetate to slow evaporation and the lid was removed for 6 hours before imaging. Humidity and temperature were measured at leaf height at ∼76% RH 22 °C for treatment and ∼95% RH 22 °C for mock. Humidity and temperature were measured using a BME280 sensor.

### Peristomatal distance measurement

Stomatal aperture is challenging to measure from confocal images, but correlates strongly with peristomatal distance ^54^, which we measured in our nlsABACUS-400n humidity treatment confocal stacks. The line tool in Fiji was used to measure distance using a transmitted-light channel.

### Foliar humidity treatment for root imaging

8 ml of ½ MS 0.8% pH 5.7 Agar was poured into a Nunc™ Lab-Tek™ II Chambered Coverglass (155360 Thermo Fisher) and allowed to set. Half of the agar was aseptically removed and seeds were placed on the agar, next to the coverslip to allow plant roots to grow vertically between the agar and coverslip (Extended Data Fig. 17). Chambers were sealed three times with micropore tape, stratified for four days and then plants were grown to 6 days post stratification in a long day chamber. For the humidity treatment, imaging chambers were opened, a piece of folded acetate was placed over the agar to prevent direct evaporation, and aerial tissues were exposed to the 40% RH 22°C chamber for six hours (Extended Data Fig. 17). Mock treatment involved opening the chamber, applying a smaller piece of acetate and resealing before returning to the growth chamber. The smaller acetate application acts as a control for any mechanical perturbation, but still retains a large area for water exchange between the agar and air, so the chamber remains humid and equilibrates quickly.

### Foliar humidity treatment for root growth assays and β- Estradiol pretreatment

80ml of ½ MS 0.8% pH 5.7 agar was poured into a 10cm square plate, and allowed to set. 2.5cm of agar was aseptically removed from one side and seeds were placed on the agar, next to the back of the plate to allow plant roots to grow vertically between the agar and plate (Extended Data Fig. 13). Plates were sealed three times with micropore tape, stratified for four days, and then plants were grown for 6 days post stratification in a long day chamber. Immediately before treatment, the position of the primary root was marked on the plate with a razor blade and a dissecting microscope.

For the humidity treatment, plates were opened, a piece of folded acetate was placed over the agar to prevent direct evaporation, and plants were exposed to the 40% RH 22°C chamber for 7 hours (Extended Data Fig. 13). Mock treatment involved opening the plates, applying a smaller piece of acetate and resealing before returning to the growth chamber. The smaller acetate application acts as a control for any mechanical perturbation, but still retains a large area for water exchange between the agar and air, so the plate remains humid and equilibrates quickly.

For UBQ10pro/SUC2pro:XVE>>CYP707A3 induction pretreatment experiments, 24 hours before humidity treatment plates were opened, sprayed with 50 μM β-Estradiol 0.25% DMSO 0.05 % Silwett-77 or mock solution (0.25% DMSO and 0.05 % Silwett-77). Excess solution was removed with a paper towel, plates were resealed and replaced in the growth chamber.

### Rootchip microfluidics treatments

The RootChip-8S device was used for ABA pulsing as previously ^12,27^. Arabidopsis seeds were germinated on the bottom 5 mm of 10 μl pipette tips filled with solidified growth medium (½ MS, 1% Agar, 0.05% (wt/vol) MES pH 5.7). After 4 to 7 d, pipette tip seedlings were transferred to the polydimethylsiloxane RootChip-8S device under aseptic conditions. A peristaltic pump was used (DNE GmbH; volumetric flow rate in each channel, 5 mL/min) to perfuse the roots with ¼ MS pH 5.7 liquid media. The dead volume was assessed, and it took approximately 12 minutes for media to pass through the tubing to reach the root, which was taken into account when plotting the ABA treatments. Imaging was performed on an inverted Leica SP8 with a 20× dry 0.70 HC PLAN APO objective. 448 nm and 514 nm lasers were used for excitation of edCerulean and edCitrine, respectively. Emission settings were 460 to 490 nm for Cerulean and 520 to 550 nm for edCitrine.

### ABA hypersensitivity germination assays

Seeds were surface sterilized, placed on large ½ MS + MES agar plates with or without 1μM ABA and stratified for 4 days. After transfer to a growth chamber, a dissecting microscope was used to score germination daily. Seedling emergence from the endosperm was used to score germination.

### ABA hypersensitivity root growth assays

Seeds were surface sterilized, placed on large ½ MS + MES agar plates vertically in a growth chamber. At 7 DAG, seedlings of approximately equal length were transferred to mock or 10 μM ABA plates. Root tip positions were marked and plates were replaced vertically in the growth cabinet for 40 hours before imaging on a flatbed scanner. Root growth was measured with the segmented line tool of Fiji.

### Confocal imaging

An upright SP8-Fliman was used for most biosensor imaging. An inverted SP8-iphox was used for RootChip imaging. All images were acquired as Z-stacks in 16 bit mode, with a 10× dry or 20× dry 0.70 HC PLAN APO dry objective. Samples were mounted in ¼ MS pH 5.7.

Typical settings were as follows: Sequential scanning was used with the following laser/detector settings: Sequence 1: 442 excitation 5-30%, HYD1: 460-500nm, 100 Gain; HYD2 525-560nm, 100 Gain. Sequence 2: 514 excitation 5-30%, HYD2 525-560nm, 100 Gain. Scan speed 400, Line averaging: 2-4, Bidirectional X:on

### Lightsheet microscope setup

Lightsheet microscopy was performed using a custom-built laser scanning light sheet microscope. The design is based on an openspim geometry^55^ with dual side illumination and dual side detection. Water immersion objectives are mounted horizontally (Nikon 10x, 0.3 NA for excitation, Olympus 20x 1.0 NA for detection) with the sample suspended from the top in an agarose filled Fluorinated Ethylene Propylene tube (FEP). For sample placement as well as for imaging the sample can be moved between the objectives well as rotated with piezo-driven stages (Nanos LPS-30, Nanos RPS-LW20). Image stacks are acquired by moving the sample through the stationary imaging plane. 445nm and 488 nm lasers (Omicron LuxX 445-100, Omicron LuxX 488-200) were used for excitation and combined in an Omicron LightHub 6 with dual fibre output. The fibre output was collimated, galvo scanned (Galvo system: Thorlabs GVSM002-EC/M) and magnified resulting in a scanned light sheet with typical FWHM < 5um. Two sCMOS cameras (Hamamatsu Orca Flash 4) with 6.5×6.5 um^2^ pixel size are used for detection. Two motorised filter wheels (Cairn OptoSpin) with bandpass filters (Semrock FF01-480/17, Semrock FF01-532/18) allow the recording of specific fluorescence bands. The microscope is controlled by a custom software developed in LabVIEW (National Instruments). Data was streamed to disk and converted to TIFF files directly after acquisition resulting in image voxel sizes of 1 μm^3^.

### Lightsheet imaging

The plants were grown suspended in a cut 10 μL pipette tip as in ^56^ in ½ MS pH5.7, 0.5% agarose FEP tubes (ID 0.8 mm)). They were illuminated from 2 sides while 3 fluorescent channels are recorded sequentially (Ch1: Exc 445nm, Em 480/17, Ch2: Exc 488nm, Em 532/18, Ch3: Exc 445nm, Em 532/18). Typical excitation powers set in software were 10%-50% for 445 nm excitation and 1-3% for 488nm excitation. Camera exposure time was set to 100ms per plane for all channels. Multiple viewpoints (60º rotation increment) were recorded for each timepoint and combined in Fiji ^57^ using the Multiview reconstruction plugin ^58^ before further analysis. Foliar ABA treatment was performed by pipetting 5 μM ABA into the top of the cut pipette tip onto the cotyledons, which is isolated from the roots.

### FRETENATOR toolset development

A fast yet flexible analysis pipeline was required to analyse biosensor data. Because the biosensors used in this paper are nuclear localized, the pipeline was designed for punctate nuclear segmentation and analysis is performed on a per nucleus basis. The toolset consists of two plugins. ***FRETENATOR Segment and ratio*** is used to segment punctate structures, perform ratio calculations and export the data as images and a results table. ***FRETENATOR ROI Labeller*** is used to assign specific labels to the regions of interest (ROI) produced by ***FRETENATOR Segment and ratio*** and exports this information to the results table.

### Development: *FRETENATOR Segment and ratio*

Fiji ^57^, an open source, multiplatform, widely adopted ImageJ^59,60^ distribution was chosen as platform to allow the greatest flexibility to users. All plug-ins were developed in jython, using CLIJ/CLIJ2 ^61^ to perform image processing directly on the graphics card. On computers with dedicated graphics cards, this allows fast analysis and modification of the segmentation settings can be performed through a graphical user interface (Extended Data Fig. 18), with near-real time segmentation previews. All code is freely available at https://github.com/JimageJ/ImageJ-Tools, along with installation and usage tutorial videos.

Segmentation steps are illustrated in Extended Data Fig. 19. Preprocessing consists of extracting the segmentation channel, applying a 3D difference of Gaussian filter to smooth noise and remove background. An optional tophat filter allows further background subtraction. A choice of various automatic methods or manual thresholding is then used to generate a binary map.

An optional 3D watershed is used to split objects. Because 3D watershed can cause the loss of too many nuclei ROI or shrink them below their original size, we compare the watershed to non-watershed binary maps. A map of the ‘lost nuclei’ is generated, which are added back later.

A 3D connected components analysis is used to generate a label map of the watershed nuclei. As a watershed shrinks objects, the labelled objects are dilated (on zero-value pixels only), then multiplied by the original threshold image. This provides a good segmentation with split objects without object shrinkage.

To correct account for any ‘lost nuclei’ absent from the image, a connected-components analysis is run on the ‘lost nuclei’ map, to generate labels which are supplemented back onto the first label map.

Once the segmentation is complete, voxels that are saturated on either the **d**onor e**x**cited **d**onor e**m**ission (DxDm), or the **d**onor e**x**cited **a**cceptor e**m**ission (DxAm) are excluded from analysis of both channels and the emission ratio (DxAm/DxDm) is calculated for each ROI. The segmentation is also used to quantify position, size, donor intensity, acceptor FRET intensity, acceptor intensity, pixel count, image frame for each ROI, which are exported as a results table along with file name, ROI identifiers (Extended Data Fig. 20). The following outputs are produced upon plugin completion: Threshold stack, the Label stack, Emission ratio stack, Emission ratio maximum Z-projection and Emission ratio nearest-point Z-projection. Please note, to halve the file size of exported images, emission ratio values are multiplied by 1000 in exported image files, allowing the files to be saved as 16-bit images, instead of 32-bit images.

A log of segmentation settings is also created every time the *FRETENATOR-Segment and ratio* plugin is run.

### Development of *FRETENATOR ROI labeller*

The ROI labeller is a follow on tool for post-segmentation analysis where users can categorize the ROI in their segmented images (Extended Data Fig. 21). It currently works on single timepoint 3D label images, allowing users to visually assign labels to one of 10 categories. Results are either output to an existing results table or can be used to remeasure a chosen image.

### *FRETENATOR* software compatibility

The majority of testing was performed on a 2017 Dell desktop (Windows 10 Intel i7-6700 CPU 3.41GHz, 32GB RAM, Intel HD Graphics 4000/AMD Radeon R7 450), and a 2014 Gigabyte laptop (Ubuntu Intel i7-4710Q 2.5GHz Quad core, 16GB RAM, Nvidia GTX 860M 4gb) on which the software runs well. We regularly use the software on various Windows, Linux and Mac machines of varying ages and specifications. Considerable speed increases are present on modern hardware with fast graphics memory. Dozens of Arabidopsis cotyledon z-stacks have been tested.

### *FRETENATOR* validation (comparison with *Imaris 8.2*)

*FRETENATOR segment and ratio* analysis was compared to the commercial software *Imaris 8*.*2* (https://imaris.oxinst.com/) for validation and to ensure comparable results. Buffer exchange and segmentation were performed as previously described ^26,62^. Segmentation in *Imaris* was performed using the surfaces wizard on the AxAm channel, with background subtraction and object splitting. The XTMeanIntensityRatio Xtension was used for emission ratio calculation.

*FRETENATOR* and IMARIS gave extremely close results in terms of both segmentation and quantification of emission ratio (Extended Data Fig. 22.). As *FRETENATOR* is free, quick to use and can be installed on old, low-specification computer hardware, *FRETENATOR* was used for subsequent biosensor analysis.

### Image analysis using *FRETENATOR*

All segmentation and labeling were performed with the *FRETENATOR* plugins. Segmentation settings were optimized for each experiment but kept constant within each experiment. The AxAm channel was used for segmentation. Watershed was used for the dense nuclei of the root tip but switched off for leaf imaging. Difference of Gaussian kernel size was determined empirically, due to different magnifications, resolutions and amount of noise. As a default, Otsu thresholds were used for segmentation, but in experiments where this gave poor segmentation, a manual threshold would be used the dataset (the same value for each image in the dataset).

For Rootchip timecourses, roots were registered in Fiji using the ‘Correct 3D drift’ plugin ^63^ before analysis.

For Lightsheet images, viewpoints were combined in Fiji ^57^ using the Multiview reconstruction plugin ^58^. Rolling ball background subtraction (Fiji: subtract background) was performed before processing with *FRETENATOR*.

### Statistical analyses and reproducibility

All statistical tests are described in the figure legends, along with sample size. Central lines indicate median and variation indicates interquartile range in box-and-whisker plots. Whiskers indicate the range. Diamonds indicate outliers.

## Acknowledgments

We would like to thank Eiji Nambara for kindly providing the *cyp707a1cyp707a3* seed and Malcolm Bennett and Poonam Mehra for providing the *snrk2*.*2snrk2*.*3, snrk2*.*2snrk2*.*3* RCH1pro::SnRK2.2 and *snrk2*.*2snrk2*.*3* SnRK2.2pro::SnRK2.2 lines and for comments on the manuscript. Thanks to Farhat Nazir, Hugo Caumon and Rui Albuquerque-Martins for help at various stages of the project. Thanks to Raymond Whiteman, Gareth Evans and Laurel Tully for their help with microscopy and horticulture. This work was funded by the Gatsby Charitable Foundation and Biotechnology and Biological Sciences Research Council (BB/P018572/1).

## Contributions

AMJ conceived of the project. AMJ, JR, MGG, RW, KS and SC designed biosensor mutations. JR, RW and MGG made DNA constructs. MGG and RW performed *in vitro* biosensor screening. JR generated plant lines, JR, MER and ML performed imaging experiments. JR performed phenotyping experiments. ML performed constructed the SPIM microscope and appropriate software. JR wrote FRETENATOR image analysis software. JR and MER performed image analysis. JR, MGG and MER analysed data and performed statistics. Protein structure prediction was performed by JR.

## Data and materials availability

New plant lines will be deposited at the Nottingham Arabidopsis Stock Centre. Binary vectors for ABACUS2 transformation as plant ABACUS2 constructs in pENTR221-f1 will be deposited at Addgene. All data has been placed Cambridge data repository. The FRETENATOR image analysis toolset, as well as installation and usage instructions are available at https://github.com/JimageJ/ImageJ-Tools.

