## Supplemental Information for "Next-generation ABACUS biosensors reveal cellular ABA dynamics driving root growth at low aerial humidity"

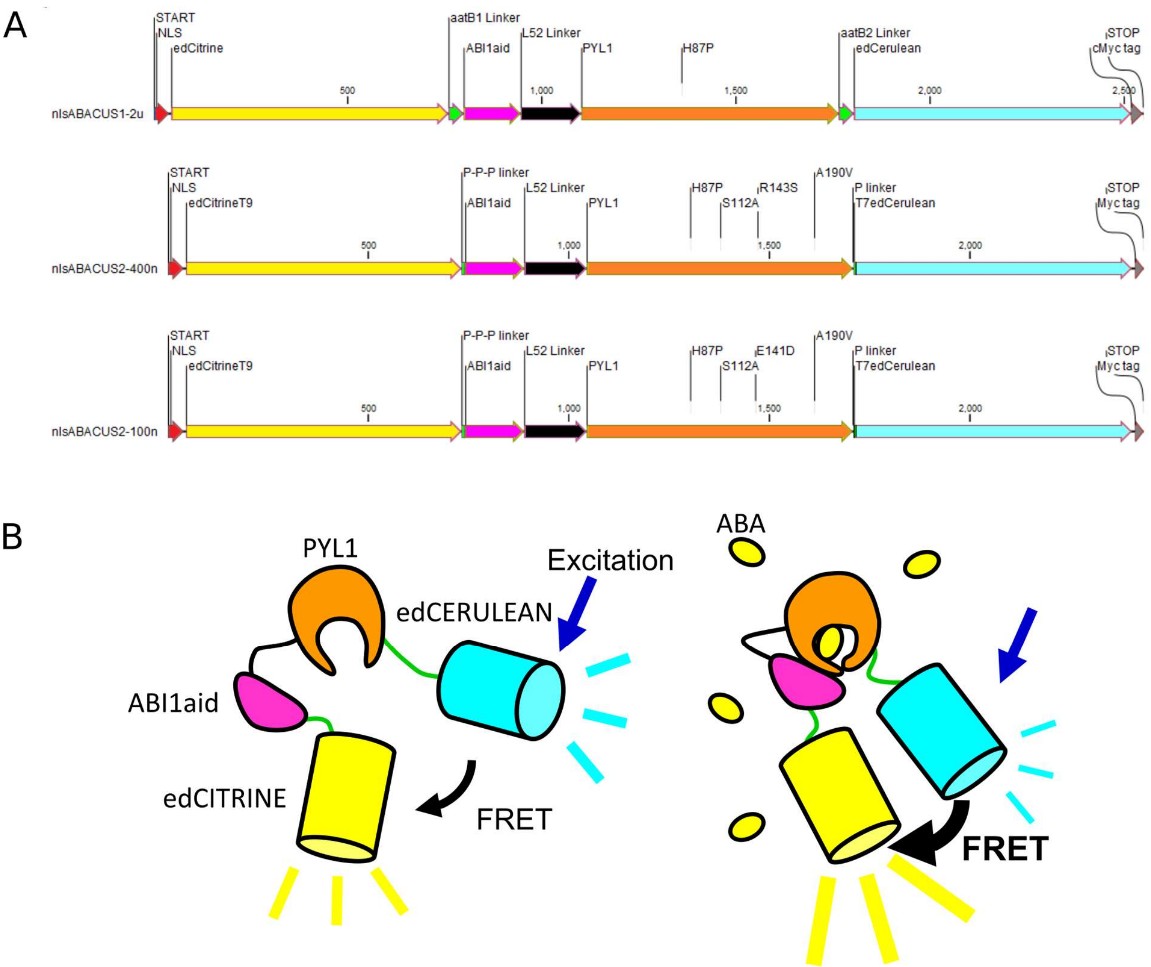


**Extended Data Fig.1. Domain and mutation annotation of ABACUS sensors**

A) Sequence annotation of nlsABACUS1-2µ, nlsABACUS2-400n (mutations: edCitrineT9, PPP-linker, PYL1: A190V, S112A, R143S; P-linker T7edCerulean) and nlsABACUS2-100n (mutations: edCitrineT9, PPP-linker, PYL1: A190V, S112A, E141D; P-linker T7edCerulean)

B) ABACUS cartoon representation in apo and holo forms. Upon ABA binding, there is a conformational change in the sensory domain (PYL1 and ABI1aid). The resultant change in fluorescent protein position means that more energy is transferred from the donor (edCERULEAN) to the acceptor (edCITRINE) via FRET.

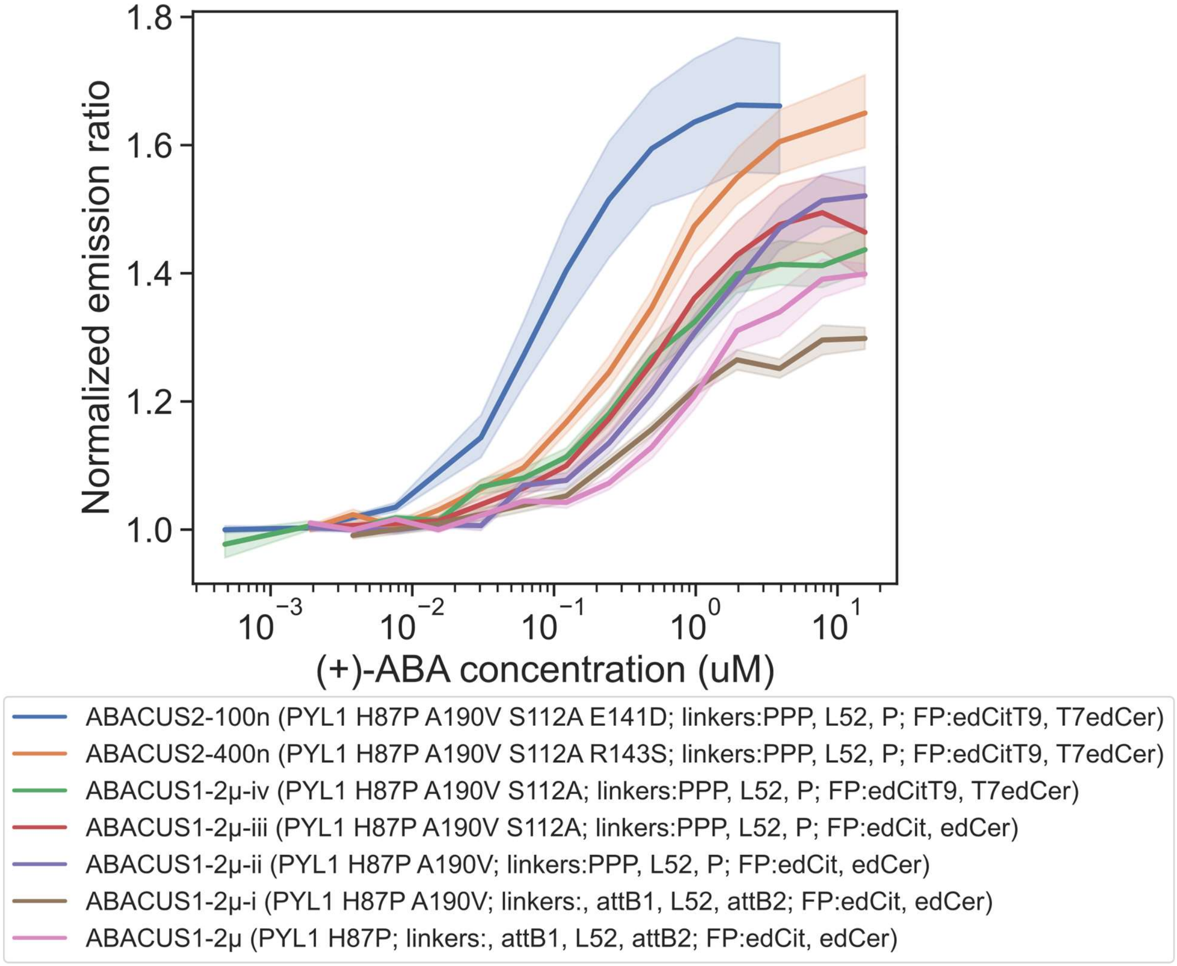


**Extended Data Fig.2. Purified ABACUS variant *in vitro* titrations.**

Yeast expressed ABACUS protein variants were purified with metal affinity chromatography and titrations were perfomed with the bioactive (+)-Abscisic acid. Emission ratio values were calculated and normalized against corresponding mock treated samples. Cit=Citrine, Cer=Cerulean. Line indicates mean of multiple independent extractions and titrations, shaded region indicates the standard error of the mean. n>3 in all cases, and are detailed in Extended Data Table 1.

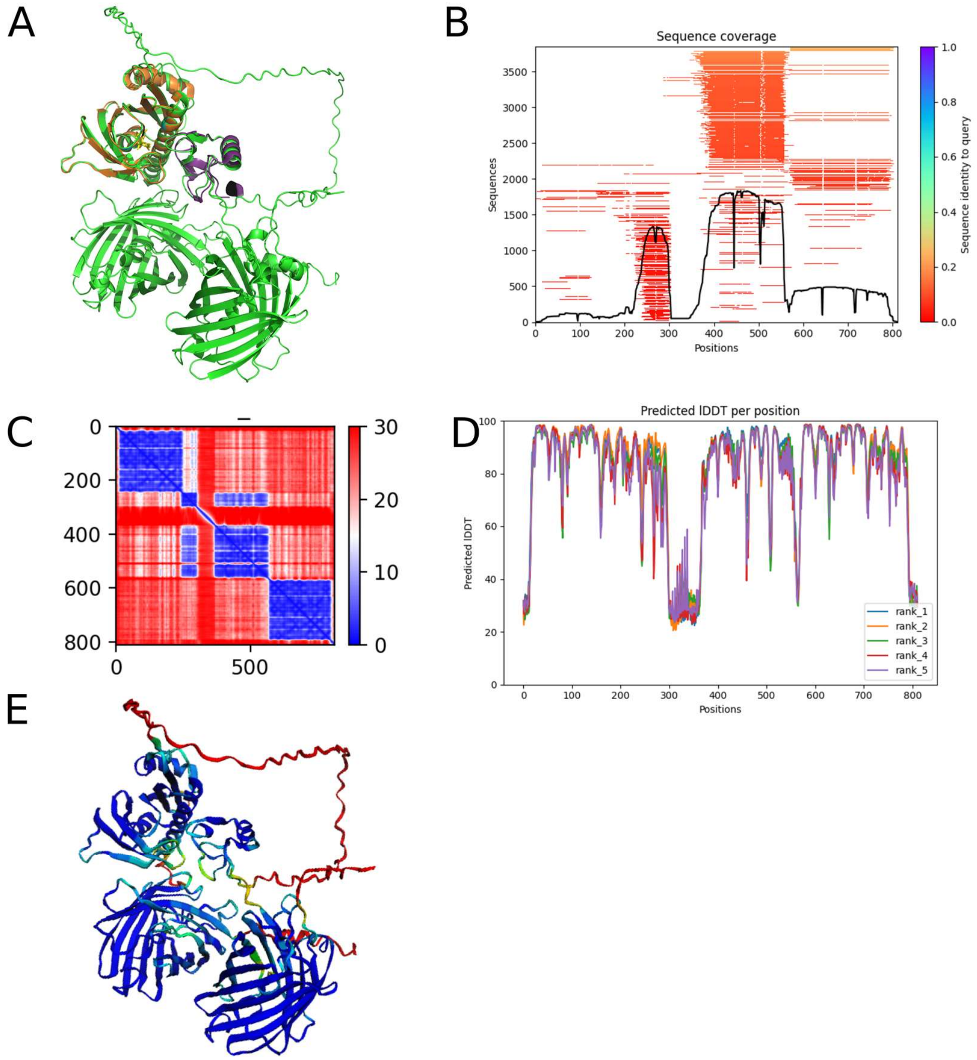


**Extended Data Fig.3. ColabFold: AlphaFold2 using MMseqs2 predictions of nlsABACUS2-100n structure**

A) Highest ranked collabfold structural prediction (green) aligned to PYL1-ABA-ABI1 structure 3JRQ. Orange: PYL1, Yellow: ABA, Magenta: ABI1 (only ABI1aid is shown). RMSD: 0.758 Å Aligned atoms: 1509, Number of refinement cycles: 5

B) Multiple sequence alignment (MSA) coverage of nlsABACUS2-100n

C) Predicted aligned error of the highest ranked structural prediction of nlsABACUS2-100n

D) pLDDT plot shows high confidence for much of the nlsABACUS2-100n structural prediction, with low confidence in the L52 spring linker

E) pLDDT mapped onto the highest ranked structural prediction. Red: low pLDDT, Blue: high pLDDT

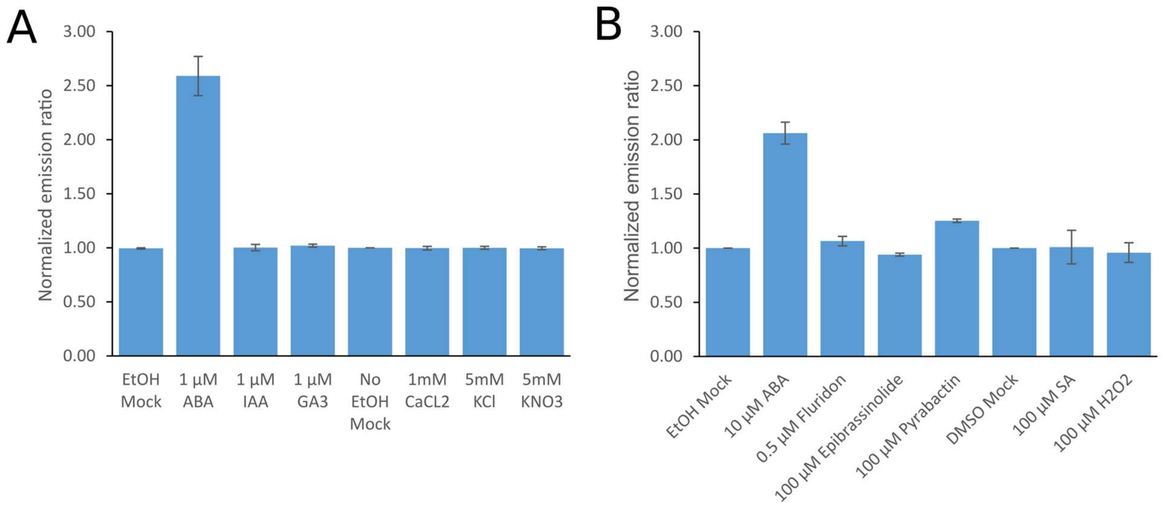


**Extended Data Fig.4. ABACUS2-100n displays high specificity to ABA and ABA signaling agonists**

A) Normalized emission ratio of purified ABACUS2-100n treated with various hormones and salts n=4. Error bars indicate standard deviation.

B) Normalized emission ratio of purified ABACUS2-100n treated with various hormones, the ABA agonist Pyrabactin and hydrogen peroxide n=2. Error bars indicate standard deviation.

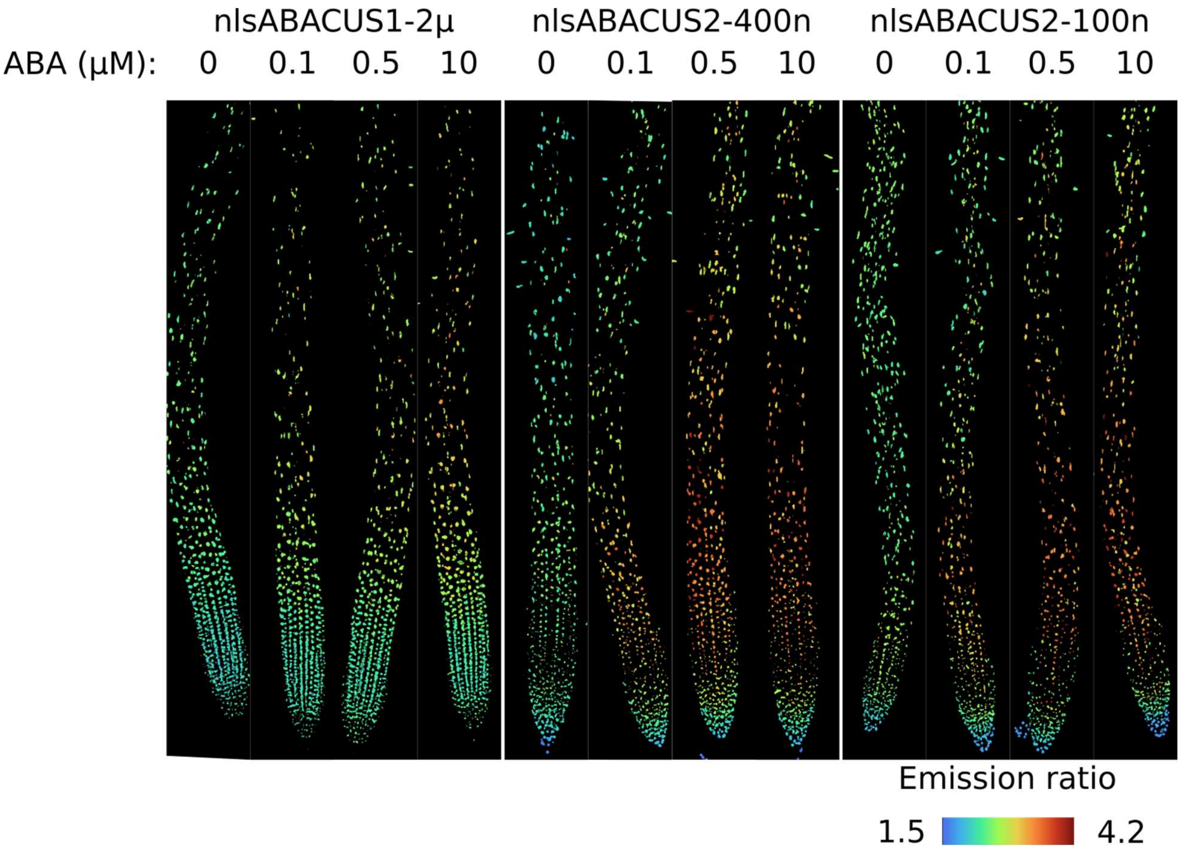


**Extended Data Fig.5. ABACUS2 responds to ABA *in planta***

Representative images nlsABACUS emission ratio responses in *Arabidopsis* roots exposed for 30 minutes to various concentrations of (±)- ABA , graphed in Fig. 1.

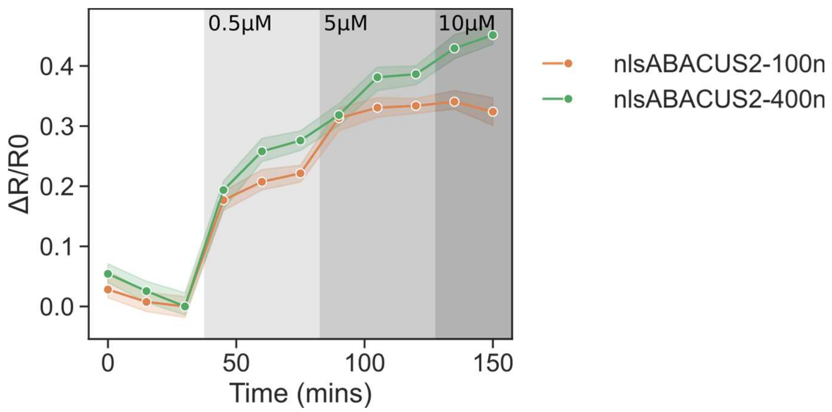


**Extended Data Fig.6. nlsABACUS2 comparison**

nlsABACUS2 sensors display fast responses to low concentrations of (±)-ABA and display a larger emission ratio change *in vivo*. ΔR/R0 is calculated as follows: (emission ratio/emission ratio immediately preceding treatment)

A

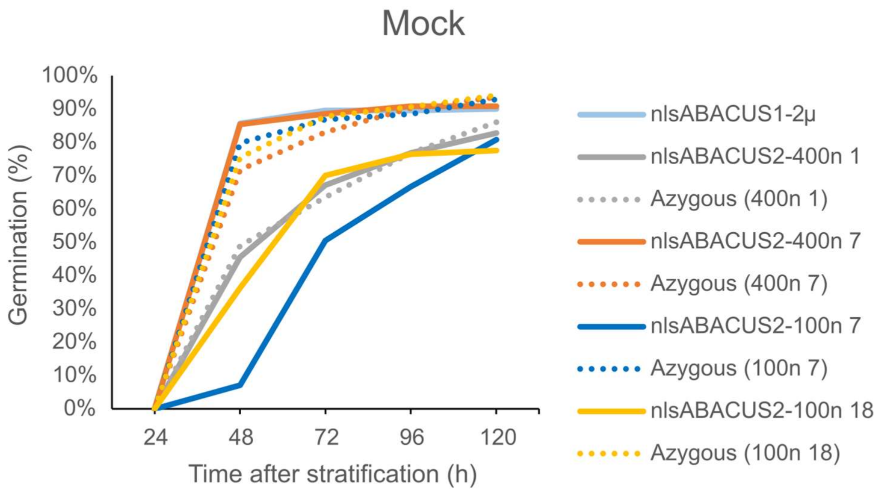


B

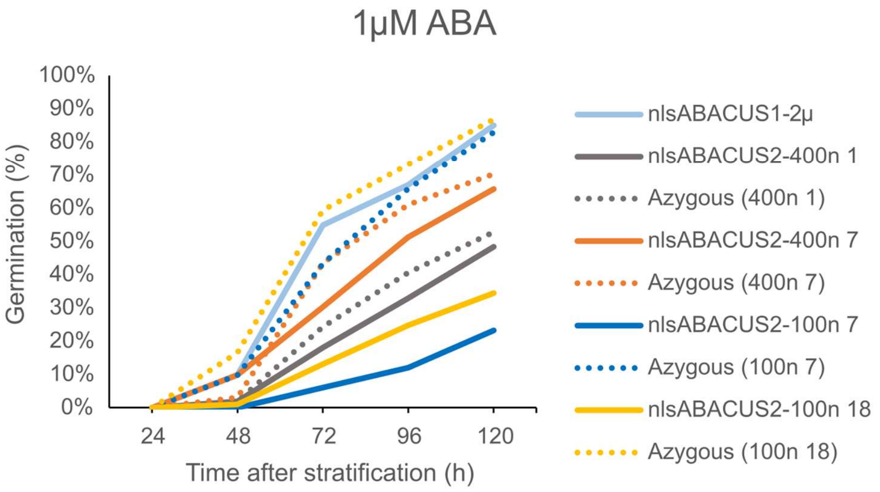


**Extended Data Fig.7. nlsABACUS2-400n and -100n display ABA hypersensitivity during germination**

A) Proportion germinated (radicle/cotyledon emergence) on 1/2MS +MES plates at 24hr intervals after placing in a growth chamber. Azygous segregants from the same transformation event were grown side by side and used as wildtype controls
B) Proportion germinated (radicle/cotyledon emergence) on 1/2MS +MES 1µM ABA plates at 24hr intervals after placing in a growth chamber. Azygous segregants from the same transformation event were grown side by side and used as wildtype controls.

n= 231 (nlsABACUS1-2µ mock), 173 (nlsABACUS1-2µ ABA),151 (nlsABACUS2-400n-1 azygous mock), 197 (nlsABACUS2-400n-1 azygous ABA), 134 (nlsABACUS2-400n-1 mock),155 (nlsABACUS2-400n-1 ABA), 124 (nlsABACUS2-400n-7 azygous mock), 131 (nlsABACUS2-400n-7 azygous ABA), 219 (nlsABACUS2-400n-7 mock), 152 (nlsABACUS2-400n-7 ABA), 183 (nlsABACUS2-100n-7 azygous mock), 164 (nlsABACUS2-100n-7 azygous ABA), 99 (nlsABACUS2-100n-7 mock), 116 (nlsABACUS2-100n-7 ABA), 204 (nlsABACUS2-100n-18 azygous mock), 175 (nlsABACUS2-100n-18 azygous ABA), 187 (nlsABACUS2-100n-18 mock), 206 (nlsABACUS2-100n-18 ABA)

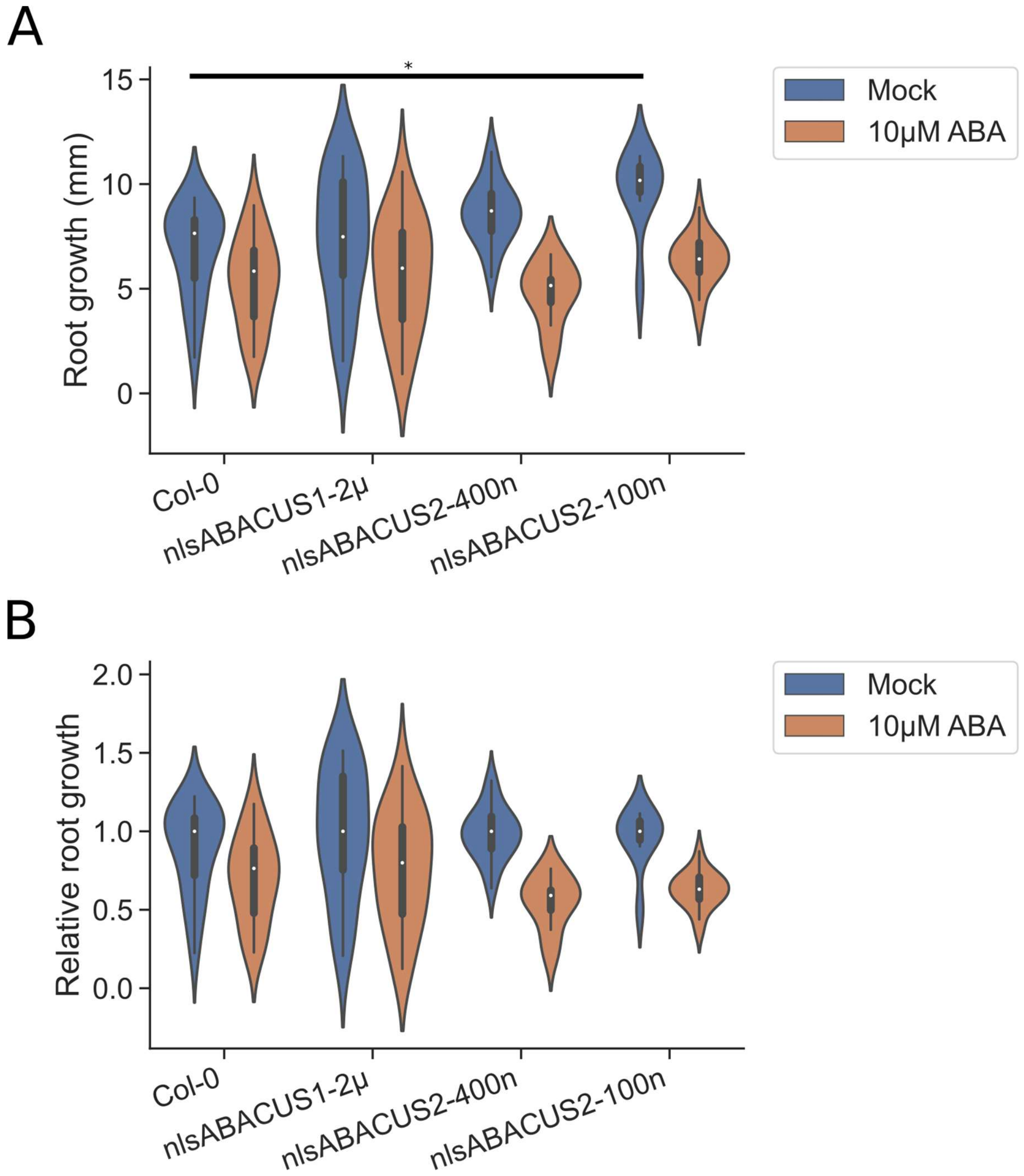


**Extended Data Fig.8. nlsABACUS2-400n and -100n display ABA hypersensitivity during root growth**

A) Root growth over 40 hours after transfer to treatment plates at 7 DAG. Two way ANOVA (Treatment: $F = 45.49$ $p<0.0001$, Genotype: $F = 4.462$, $p=0.0051$, Interaction: $F = 2.570$, $p=0.0572$). A Tukey post hoc test was used for multiple comparisons. Asterisks indicate statistical significance *:$p<0.05$, **:$p<0.01$, ***:$p<0.001$, ****:$p<0.0001$

B) Root growth normalized to the median root growth over 40 hours after transfer to treatment plates at 7 DAG.

n=19,17,19,20,19,13,9,19 respectively

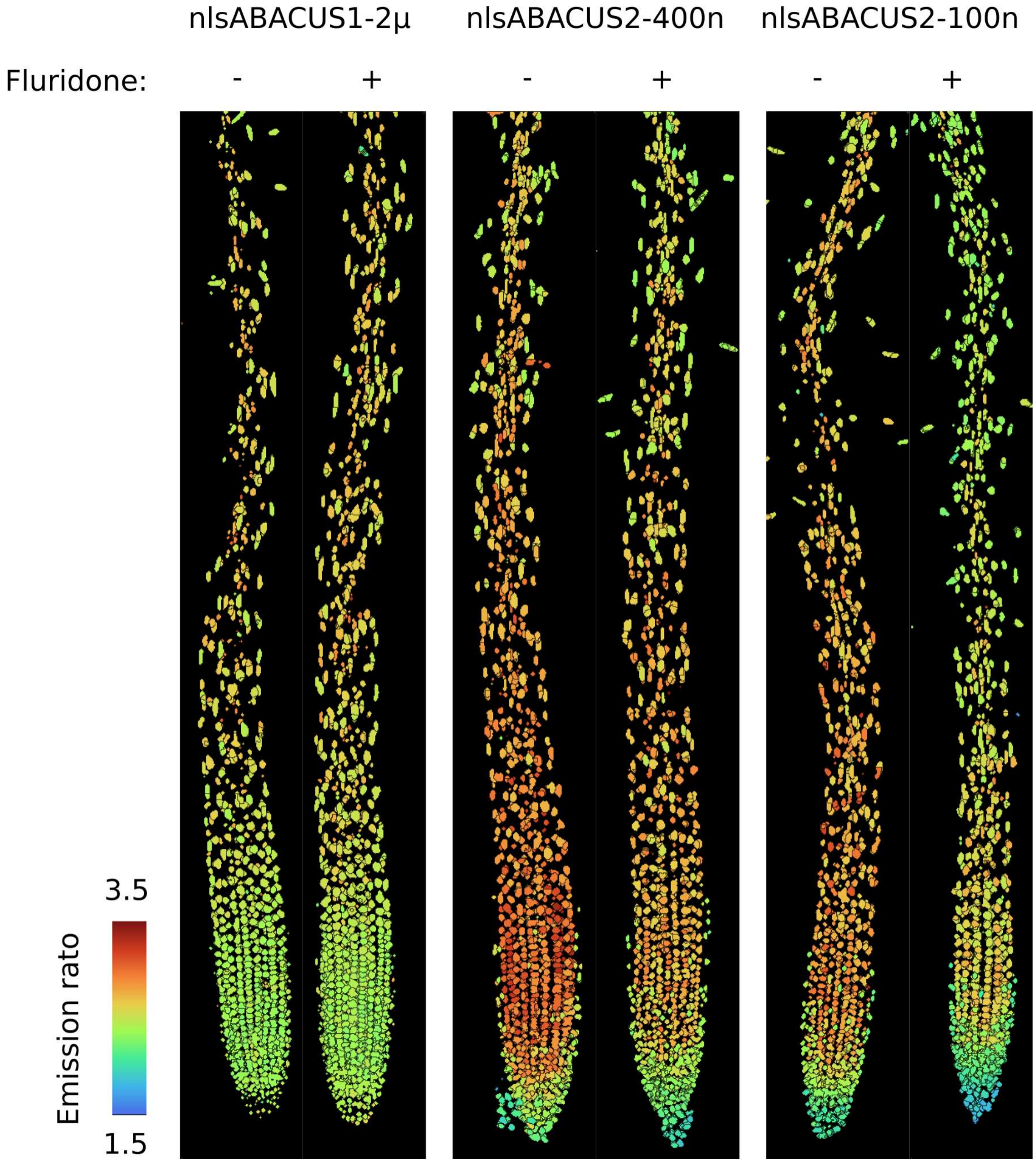


**Extended Data Fig.9. Inhibiting ABA biosynthesis reduces nlsABACUS2 emission ratios**

Representative images of emission ratios in nlsABACUS expressing Arabidopsis roots after 24hr of 0.4 µM fluridone or mock treatment.

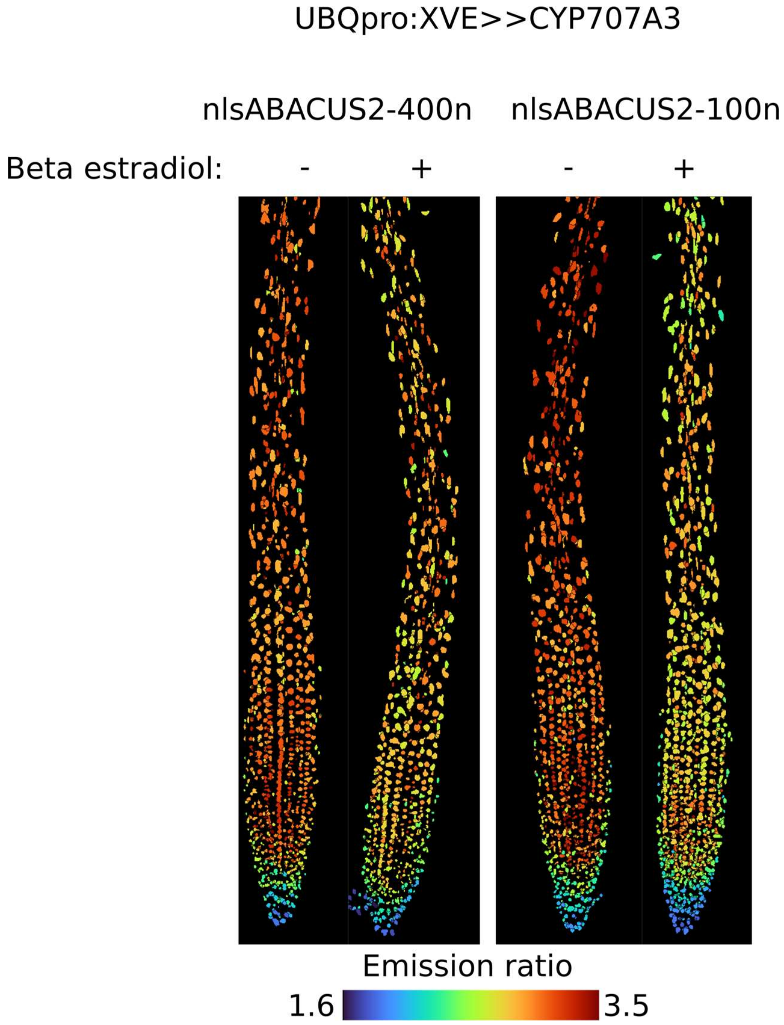


**Extended Data Fig.10. Inducing ABA catabolism reduces nlsABACUS2 emission ratios**
Representative images of emission ratios in nlsABACUS2 expressing roots after 24hr mock or 10uM β-estradiol UBQ10pro:XVE>>CYP707A3 induction. Corresponds to graph of multiple roots in Fig. 2D.

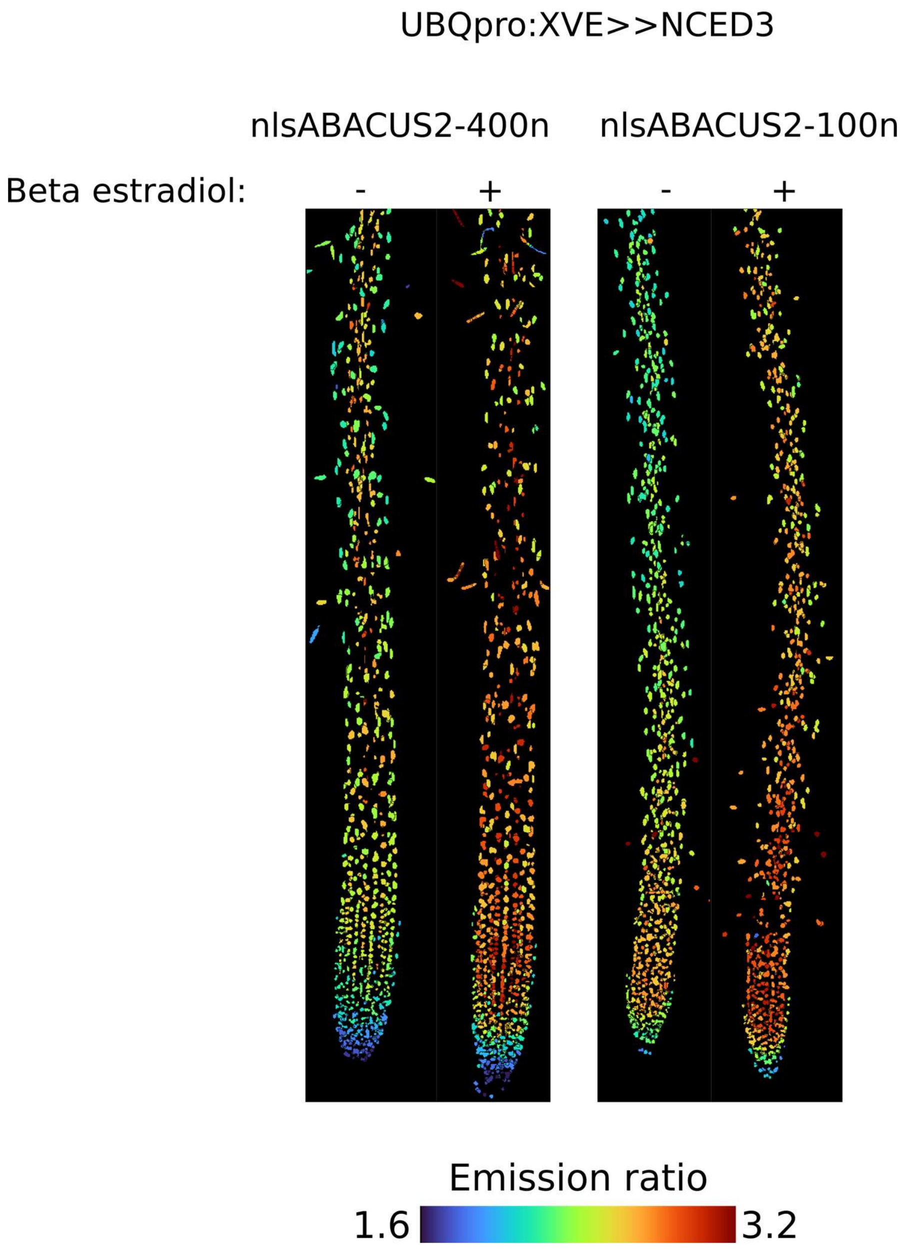


**Extended Data Fig.11. Inducing ABA biosynthesis increases nlsABACUS2 emission ratios**

Representative images of emission ratio changes in nlsABACUS2 expressing roots after 24hr mock or 10 μM β-estradiol UBQ10pro:XVE>>NCED3 induction. Corresponds to graph of multiple roots in Fig. 2C.

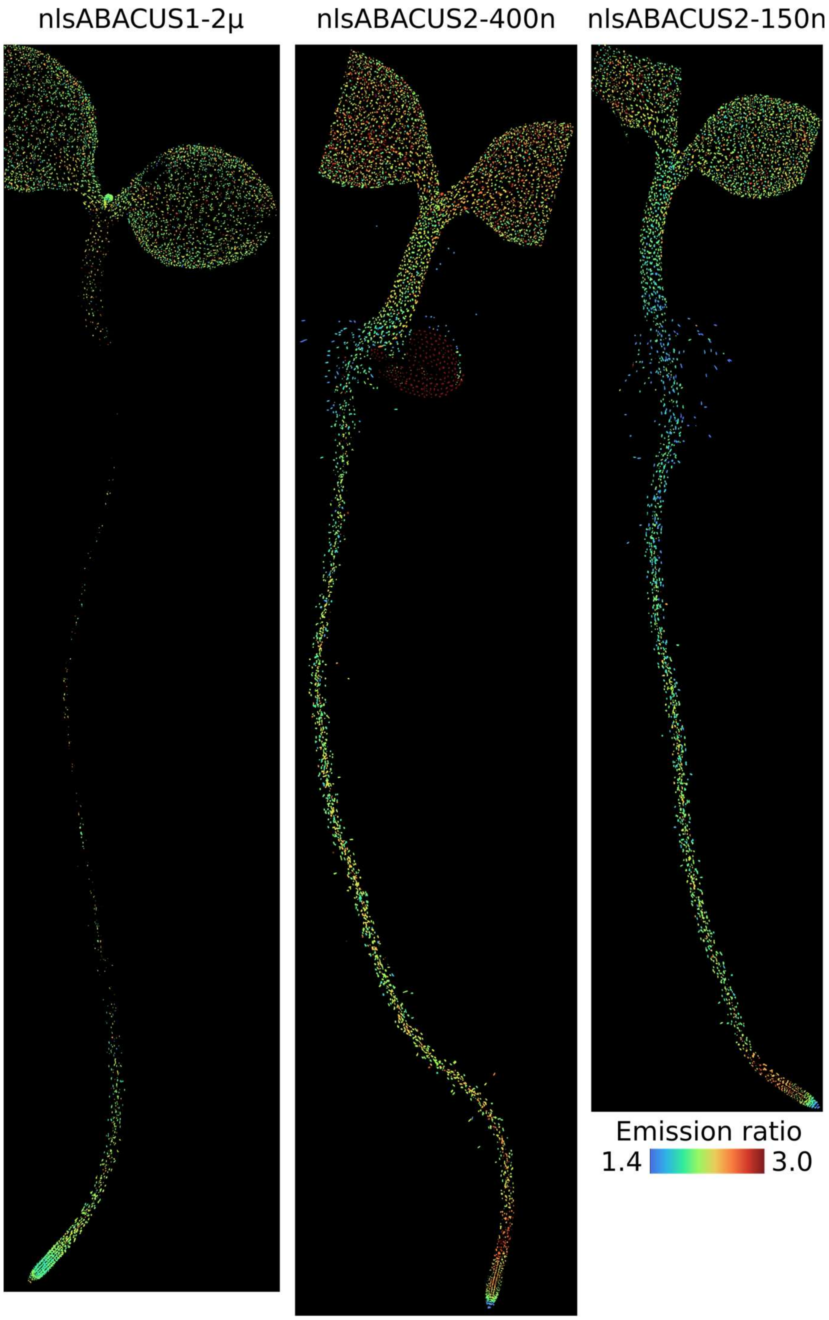


**Extended Data Fig.12. nlsABACUS2s can be used to map ABA patterns *in planta***
nlsABACUS2 expressing *Arabidopsis* seedlings display high emission ratios in the internal tissues of the cotyledons, the vasculature and the root elongation zone/meristem/differentiation zones.

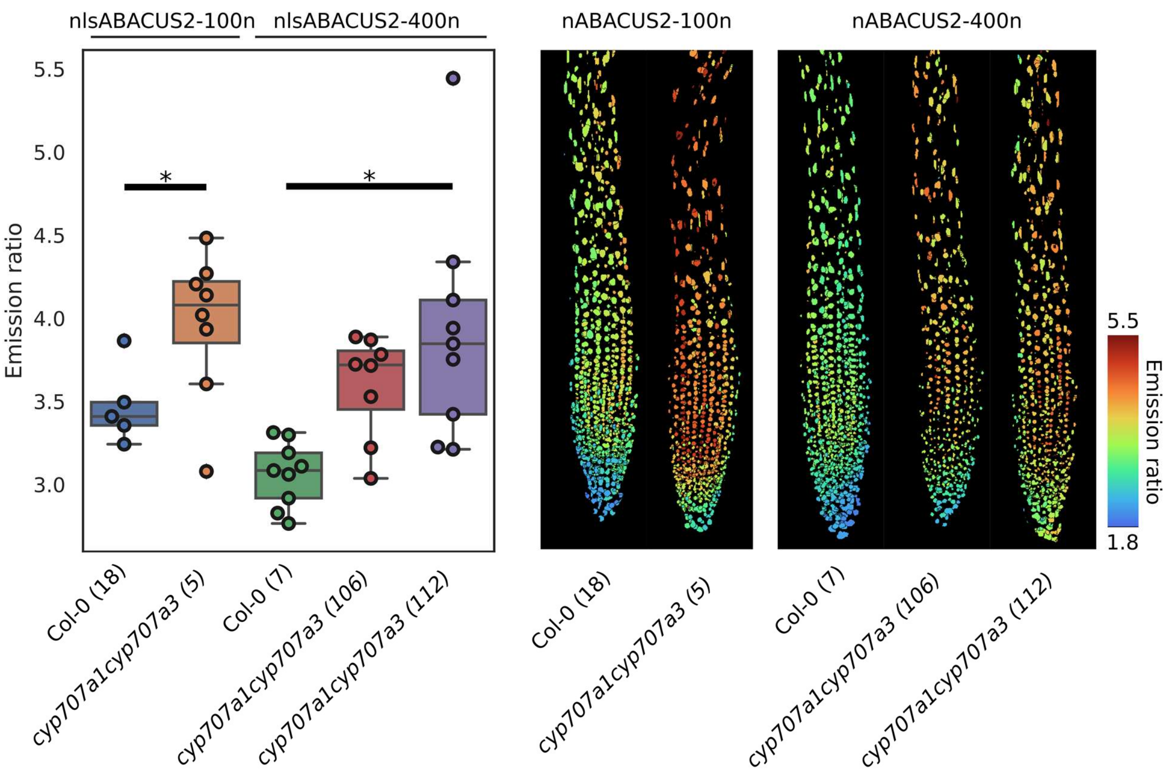


**Extended Data Fig.13. nlsABACUS2s demonstrate that *cyp707a1cyp707a3* hyperaccumulate ABA in roots**

nlsABACUS2-400n and nlsABACUS2-100n plants both display higher emission ratios in the catabolism double mutant *cyp707a1cyp707a3* implying that these genes are essential in unstressed conditions to prevent ABA overaccumulation. For nlsABACUS2-400n lines asterisks indicate statistical significance with a Tukey multiple comparison test within sensor group. For nlsABACUS2-100n lines unpaired two tail T test (t=2.275 df=11 p=0.0439), *:p<0.05, **:p<0.01, ***:p<0.001, ****:p<0.0001.
n=5,8,9,8,9 respectively

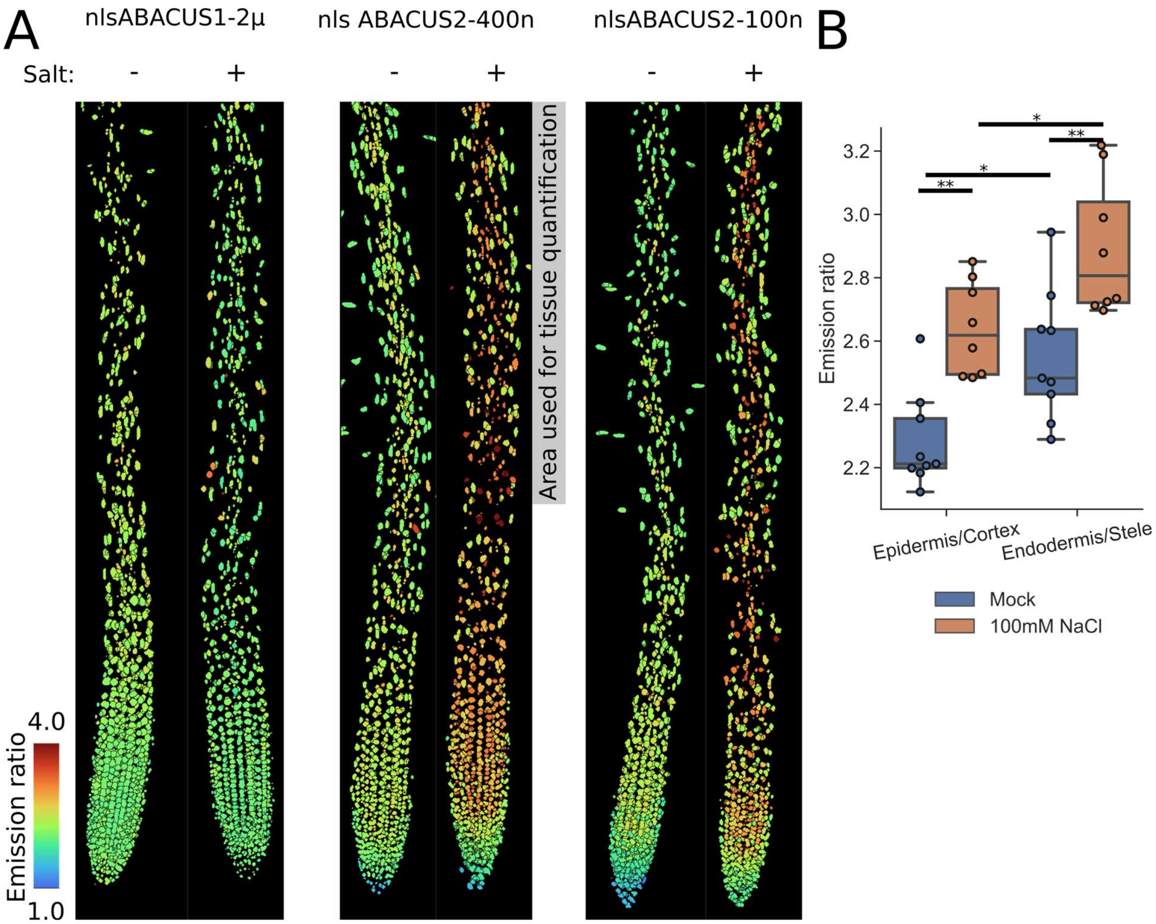


**Extended Data Fig.14. The central axis (endodermis and stele) accumulates more ABA than the epidermis and cortex under unstressed and salt conditions**

A) ABACUS2 shows increased emission ratios following 5.5 hours of 100mM NaCl treatment (Fig 3f reproduced at higher resolution to enable tissue types to be seen). The zone used for quantification of radial ABA accumulation indicated.

B) nlsABACUS2-400n emission ratios are higher in the endodermis/stele than the epidermis/cortex in the absence and presence of 100mM NaCl. Two way ANOVA (Tissue: F=17.38 DF=1, P=0.0002, Treatment: F=30.74 DF=1 p<0.0001, Interaction F=0.0199, DF=1, P=0.889) A Tukey post hoc test was used for multiple comparisons. Asterisks indicate statistical significance *:p<0.05, **:p<0.01, ***:p<0.001, ****:p<0.0001. n=9,8,9,8 respectively

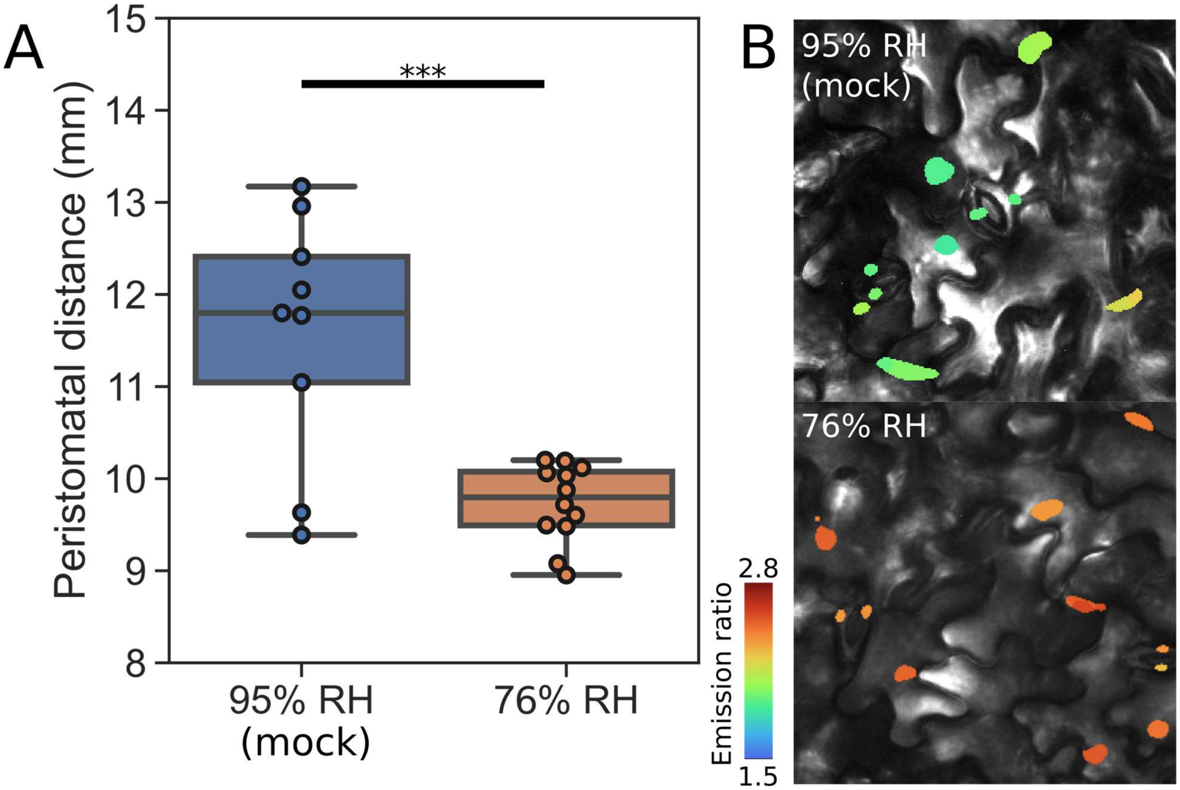


**Extended Data Fig.15. Decreasing foliar humidity increases leaf epidermal ABA levels and closes stomata**

A) Peristomatal distance for 15 DAG nlsABACUS2-400n plants decreases following a 6-hour humidity reduction. Datapoints indicated the mean of distance of all stomata in a field of view of a confocal datastack. Each field of view contained between 8 and 16 measured stomata. Peristomatal distance taken to be a proxy for stomatal aperture [54]. Unpaired two tailed T-test p=0.0002, t=4.501, df=19. n=9,12 respectively

B) Cropped representative images of nlsABACUS2-400n emission ratios in response to a 6-hour humidity decrease. Relative humidity (RH) indicates the measured humidity at leaf height during the treatments.

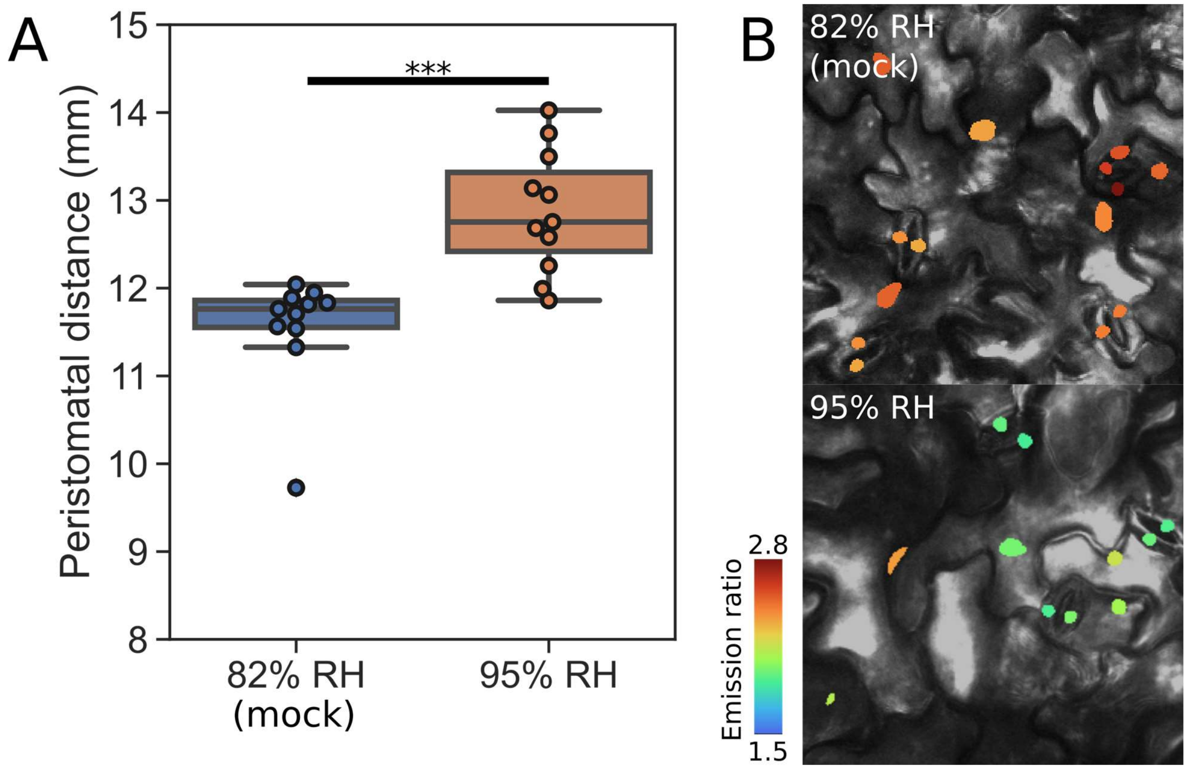


**Extended Data Fig.16. Increasing foliar humidity decreases leaf epidermal ABA levels and opens stomata**

A) Peristomatal distance for 15 DAG nlsABACUS2-400n plants decreases following a 6-hour humidity increase. Datapoints indicated the mean of distance of all stomata in a field of view of a confocal datastack. Each field of view contained at between 11 and 20 measured stomata. Peristomatal distance taken to be a proxy for stomatal aperture [54]. Unpaired two tailed T-test $p=0.0002$, $t=4.584$, $df=20$. $n=10,11$ respectively

B) Cropped representative images of nlsABACUS2-400n emission ratios in response to a 6-hour humidity increase. Relative humidity (RH) indicates the measured humidity at leaf height during the treatments.

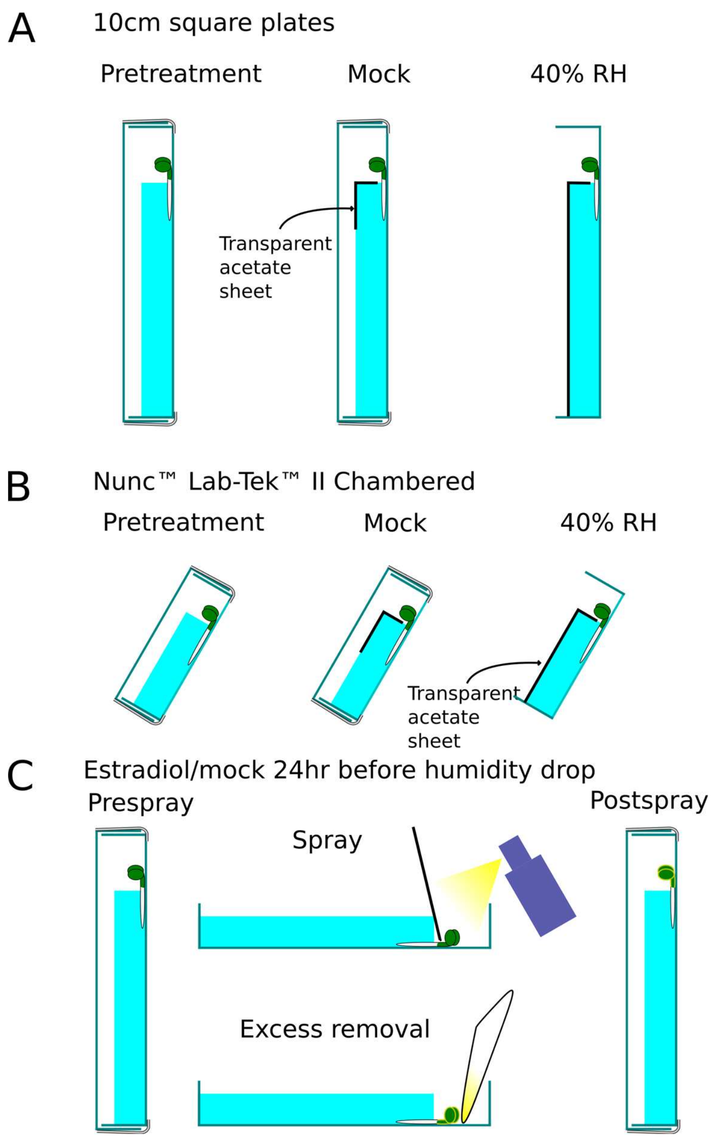


**Extended Data Fig.17. Aerial humidity treatments with hydrated roots for root growth assays and imaging assays.**

A) Root growth assay where plates are opened and a piece of folded acetate sheet (thick black line) is used to cover all or part of the agar (blue). Mock plates are resealed and all plates are placed back in a growth chamber set to 40% RH.
B) Root imaging assay, where chambered coverglasses are opened and a piece of folded acetate sheet is used to cover all or part of the agar. Mock chambered coverglasses are resealed and all chambered coverglasses are placed back in a growth chamber set to 40% RH.
C) 50 µM β-estradiol treatment regime 24hr before humidity treatment for root growth assays.

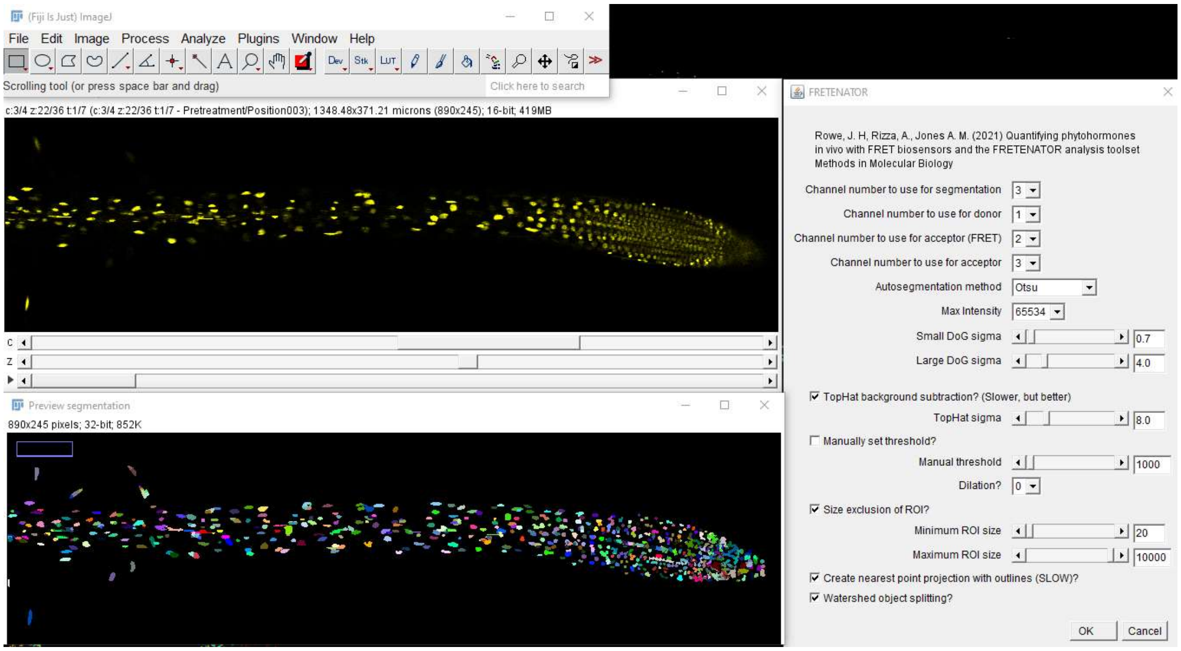


**Extended Data Fig.18. *FRETENATOR segment and ratio* user interface**
The user interface of FRETENATOR-Segment and ratio FIJI plugin. Various dropdowns, sliders and checkboxes on the plugin window (right) can be used to alter the segmentation settings, which will update the segmentation preview (bottom left). The Plugin window also allows channel selection for ratio calculations.

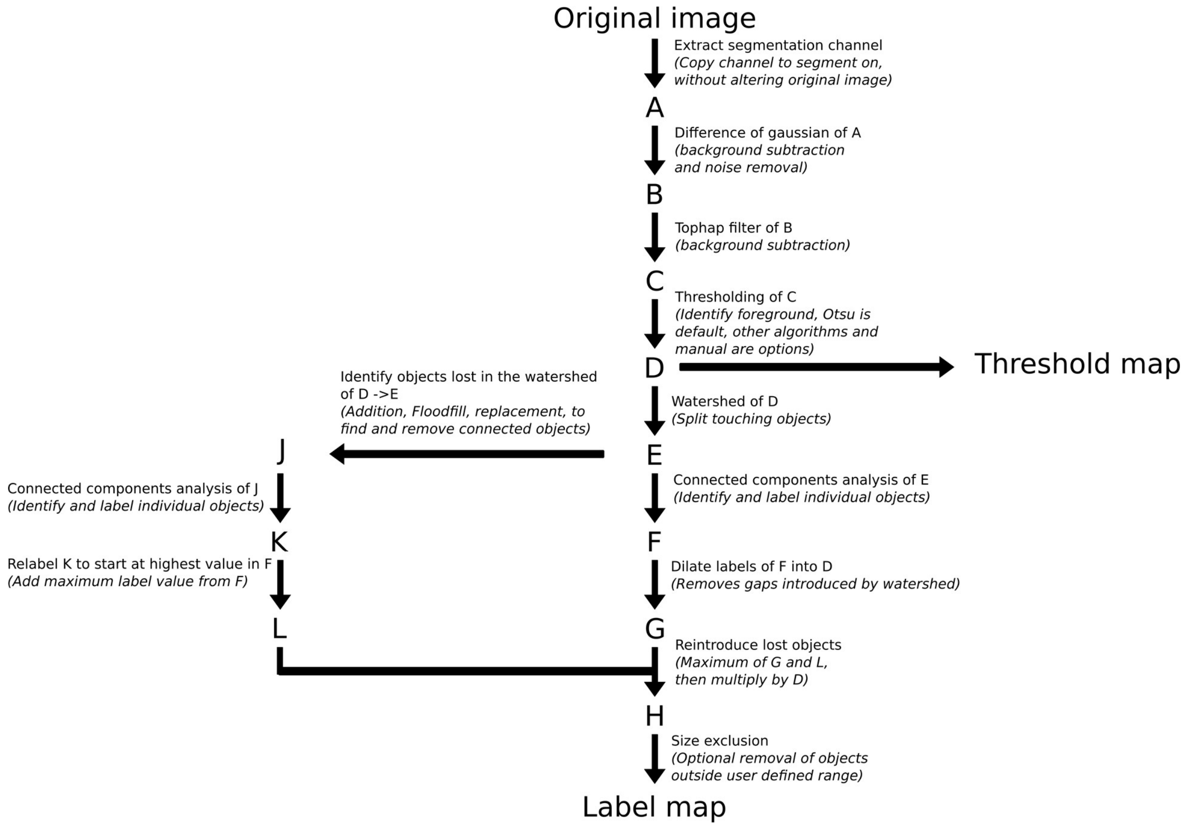


**Extended Data Fig.19.** ***FRETENATOR segment and ratio*** **segmentation pipeline**
Segmentation steps used in the FRETENATOR-Segment and ratio FIJI plugin.

Letters indicate intermediate images used in downstream processing. Arrows indicate processing steps.

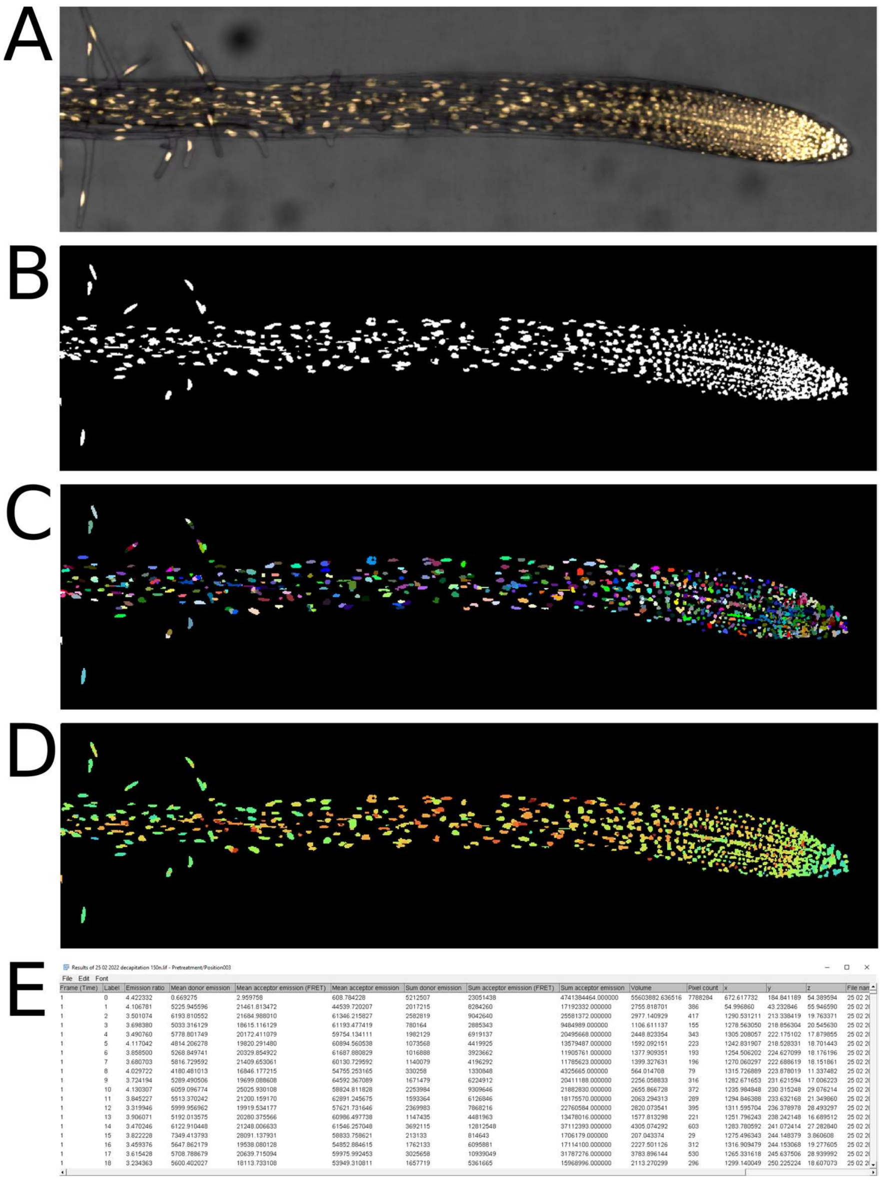


**Extended Data Fig.20. Original image and *FRETENATOR-Segment and ratio* outputs**

A) Typical import image stack, with at least two channels to calculate ratios.

B) 3D/4D stack of the thresholded image *FRETENATOR* output where foreground is assigned 1 and background is assigned 0

C) 3D/4D 'Label map' *FRETENATOR* output where each nucleus is given a unique identifier value (Displayed with the Glasbey on Dark LUT)

D) 3D/4D emission ratio map *FRETENATOR* output where each nucleus is assigned the calculated emission ratio X1000. *FRETENATOR* will also output a maximum Z-projection and outlined nearest point Z-projection of the emission ratio stack. NB: To halve the file size of exported images, emission ratio values are multiplied by 1000 in exported image files, allowing the files to be saved as 16-bit images (instead of 32-bit float images).

E) A new results table *FRETENATOR* output which can be saved as a .csv. This details the measurements of each nucleus (centroid position, size, DxDm, DxAm, AxAm, pixel count, image frame, file name, ROI identifiers)

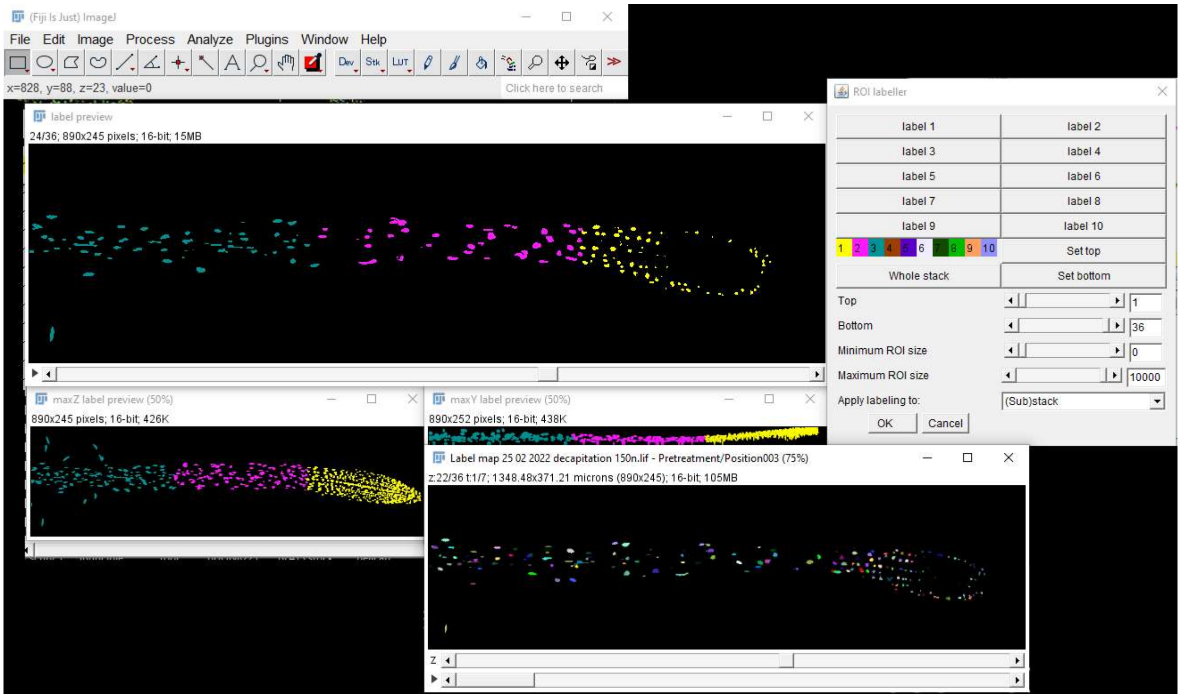


**Extended Data Fig.21. User interface of *FRETENATOR-ROI labeller***

ImageJ selection tools (e.g. box, ellipses, line etc.) can be used to select parts of the label preview (top left) and then assign them specific labels by clicking the a button on the user plugin menu panel (right). This will automatically update the preview windows.

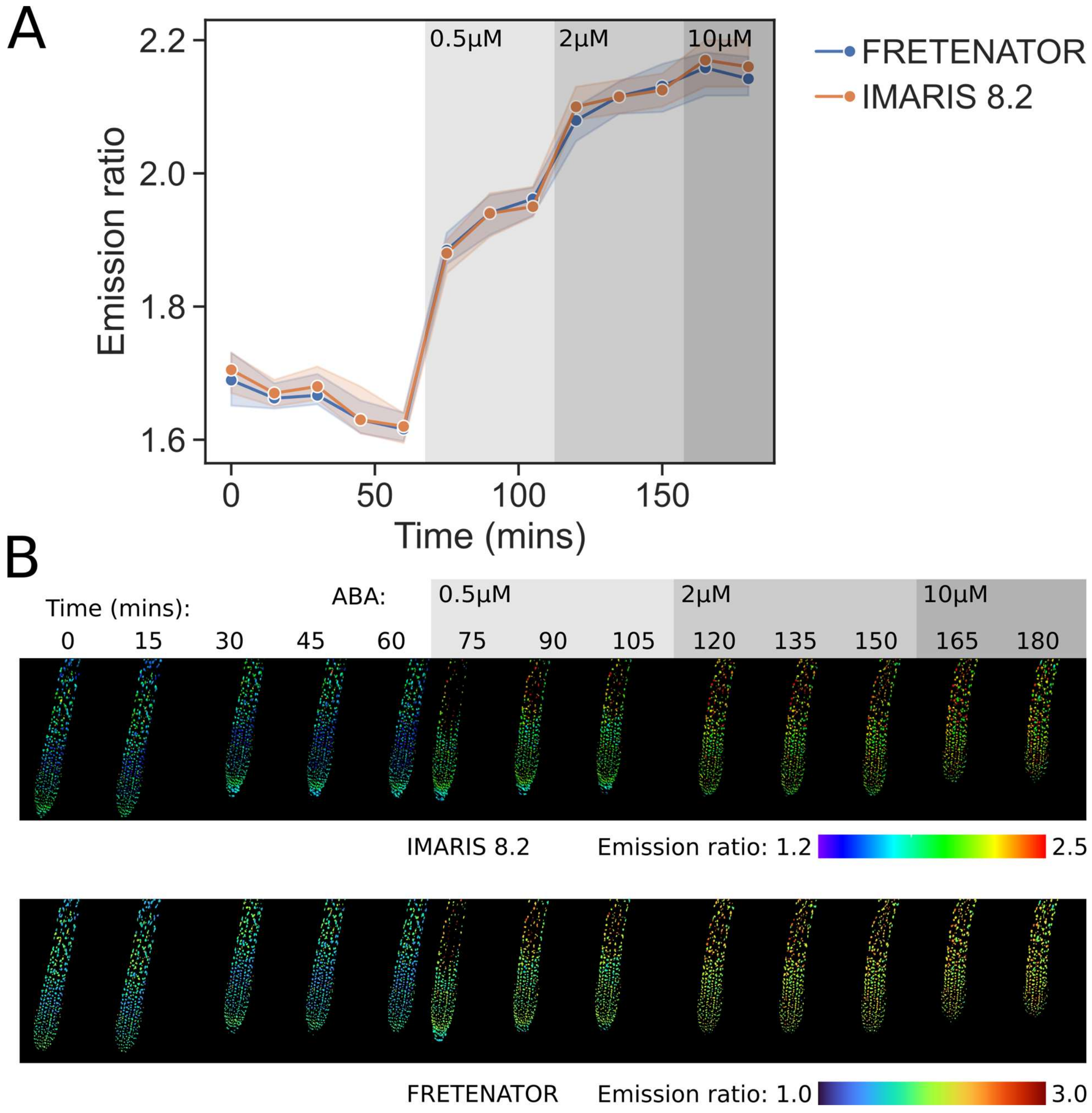


**Extended Data Fig.22. FRETENATOR and Imaris 8.2 give similar segmentation and results**

A) Quantification of emission ratios of FRETENATOR and IMARIS 8.2 shows the same trend when quantifying an nlsABACUS2-100n ABA treatment. Line indicates median

B) FRETENATOR and IMARIS 8.2 show similar segmentation and emission ratio patterns when quantifying an nlsABACUS2-100n ABA treatment.

Note this experiment is a software validation, and so uses the same dataset as Extended Data Fig.4.

**Extended Data Table 1. Emission ratios of purified ABACUS variants *in vitro*.**

| Sensor | AVERAGE $K_D$ | SD $K_D$ | AVERAGE RC (%) | SD RC | n |
|---|---|---|---|---|---|
| **ABACUS1-2µ**<br>PYL1 H87P<br>Linkers: attB1-L52-attB2<br>FP: edCit, edCer | **1.143** | **0.499** | **43.0** | **5.0** | **6** |
| ABACUS1-2µ-i<br>PYL1 H87P, A190V<br>Linkers: attB1-L52-attB2<br>FP: edCit, edCer | 0.911 | 0.047 | 30.3 | 4.7 | 3 |
| ABACUS1-2µ-ii<br>PYL1 H87P, A190V<br>Linkers: PPP-L52-P<br>FP: edCit, edCer | 0.948 | 0.460 | 63.3 | 14.7 | 12 |
| ABACUS1-2µ -iii<br>PYL1 H87P, A190V, S112A<br>Linkers: PPP-L52-P<br>FP: edCit, edCer | 0.578 | 0.300 | 68.9 | 15.5 | 5 |
| ABACUS1-2µ-iv<br>PYL1 H87P, A190V, S112A<br>Linkers: PPP-L52-P;<br>FP: edCitT9, T7edCer | 0.472 | 0.231 | 52.3 | 10.4 | 15 |
| **ABACUS2-400n**<br>PYL1 H87P, A190V, S112A, R143S<br>Linkers: PPP-L52-P<br>FP: edCitT9, T7edCer | **0.445** | **0.222** | **71.0** | **19.2** | **16** |
| **ABACUS2-100n**<br>PYL1 H87P, A190V, S112A, E141D<br>Linkers: PPP-L52-P<br>FP: edCitT9, T7edCer | **0.098** | **0.037** | **67.6** | **25.8** | **13** |

**Extended Data Table 2. Primers used in this study**

SDM primers for ABACUS variants:

| QuikChange Primer | Sequence |
|---|---|
| PYL1 A190V Fw | gattcaatctaataaccgtatcaacaaaacaatctcgtatcttcctcc |
| PYL1 A190V Rv | ggaggaagatacgagttgtttgttgatacggttatagattgaatc |
| PYL1 S112A Fw | ccggtaatccagctatcacgttcacgtcgcgcgt |
| PYL1 S112A Rv | acgcgcgacgtgaacgtgatagctggattaccgg |
| PYL1 E141D Fw | gtttagtataaccggtggtgatcataggctgaggaattataaa |
| PYL1 E141D Rv | tttataattcctcagcctatgatcaccaccggttatactaaac |
| PYL1 R143S Fw | accggtggtgaacatagcctgaggaattataaatcg |
| PYL1 R143S Rv | cgatttataattcctcaggctatgttcaccaccggt |

Linker swap, fluorescent protein truncation and citrine codon diversification primers:

| In-Fusion Primer | Sequence |
|---|---|
| Cit 3' ABI1aid 5' Fwd | ggacgagctgtacaagcataaaccggatagagaagatga |
| edCit 3' plus 1 proline ABI1aid 5' Fwd | ggacgagctgtacaagccacataaaccggatagagaagatga |
| edCit 3' plus 2 prolines ABI1aid 5' Fwd | ggacgagctgtacaagccacctcataaaccggatagagaagatga |
| edCit 3' plus 3 prolines ABI1aid 5' Fwd | ggacgagctgtacaagccacctccacataaaccggatagagaagatga |
| edCer 5' Rev PYL1 3' Rev | cctcgcccttgctcaccctaacctgagaagagttgttgt |
| edCer 5' Rev plus 1 proline PYL1 3' Rev | cctcgcccttgctcacaggcctaacctgagaagagttgttgt |
| edCer 5' Rev plus 2 prolines PYL1 3' Rev | cctcgcccttgctcacaggtggcctaacctgagaagagttgttgt |
| edCer 5' Rev plus 3 prolines PYL1 3' Rev | cctcgcccttgctcacaggtggaggcctaacctgagaagagttgttgt |
| edCit Codon Diversified-T9 fw | ccacctccacataaaccgga |
| edCit Codon Diversified-T9 rv | tttatgtggaggtggtataccagcagcagt |
| T7-edCer fw | tctcaggttaggcctttcaccggggtggtg |
| T7-edCer rv | aggcctaacctgagaagagt |

Primers to generate gateway constructs and replace codon diversified citrine:

| | Name | Sequence | Amplified with | Description |
|---|---|---|---|---|
| 1 | NCED3 CDS F1 attB1 | GGGG ACA AGT TTG TAC AAA AAA GCA GGC TTT atg gct tct ttc acg gca ac | 2 | Multisite gateway |
| 2 | NCED3 CDS R1 attB2 | GGGG AC CAC TTT GTA CAA GAA AGC TGG GT tca cac gac ctg ctt cg | 1 | Multisite gateway |
| 3 | pDONR backbone F | GATTTATAATAGACCCAGCTTTCTTGTACAAAG | 4 | infusion cloning of ABACUS2 non codon diversified edCitrine |
| 4 | pDONR backbone NLS R | ACCTCGAGCCCTCCAACCTTTCTCTTCTTC | 3 | infusion cloning of ABACUS2 non codon diversified edCitrine |
| 5 | Start NLSedCit F | TTGGAGGGCTCGAGGTGAGCAAGGG | 6 | infusion cloning of ABACUS2 non codon diversified edCitrine |
| 6 | edCIT R | GGAGGTGGCTTGTACAGCTCGTCCATGC | 5 | infusion cloning of ABACUS2 non codon diversified edCitrine |
| 7 | ABI1aid F | TGTACAAGCCACCTCCACATAAACCGGATA | 8 | infusion cloning of ABACUS2 non codon diversified edCitrine |
| 8 | myc STOP R | CTGGGTCTATTATAAATCTTCCTCACTTATC | 7 | infusion cloning of ABACUS2 non codon diversified edCitrine |

**Extended Data Table 3.** ***Arabidopsis*** **germplasm used in this study**

| Line | Background | Creation method | Origin | Selection | NASC ID |
|---|---|---|---|---|---|
| Col-0 | | | | | |
| UBQ10pro::nlsABACUS2-400n line 1 | Col-0 | Floral dip | This study | FAST-RED | TBD |
| UBQ10pro::nlsABACUS2-400n line 7 | Col-0 | Floral dip | This study | FAST-RED | TBD |
| UBQ10pro::nlsABACUS2-100n line 7 | Col-0 | Floral dip | This study | FAST-RED | TBD |
| UBQ10pro::nlsABACUS2-100n line 18 | Col-0 | Floral dip | This study | FAST-RED | TBD |
| P16pro::nlsABACUS1-2µ | Col-0 | Floral dip | This study | | TBD |
| UBQ10pro:XVE>>NCED3 line 33 | Col-0 | Floral dip | This study | Hygromycin | TBD |
| UBQ10pro:XVE>>CYP707A3 line 99 | Col-0 | Floral dip | This study | Hygromycin | TBD |
| SUC2pro:XVE>> CYP707A3 line 194 | Col-0 | Floral dip | This study | Hygromycin | TBD |
| UBQ10pro:XVE>>NCED3 UBQ10pro::nlsABACUS2-100n line 1 | UBQ10pro:XVE>>NCED3 line 33 | Floral dip | This study | Hygromycin, FAST-RED | TBD |
| UBQ10pro:XVE>>NCED3 UBQ10pro::nlsABACUS2-400n line 6 | UBQ10pro:XVE>>NCED3 line 33 | Floral dip | This study | Hygromycin, FAST-RED | TBD |
| UBQ10pro:XVE>>CYP707A3 UBQ10pro::nlsABACUS2-100n line 1 | UBQ10pro:XVE>>CYP707A3 line 99 | Floral dip | This study | Hygromycin, FAST-RED | TBD |
| UBQ10pro:XVE>> CYP707A3 UBQ10pro::nlsABACUS2-400n line 2 | UBQ10pro:XVE>>CYP707A3 line 99 | Floral dip | This study | Hygromycin, FAST-RED | TBD |
| *snrk2.2snrk2.3* | Col-0 | - | [43] | - | |
| *snrk2.2snrk2.3* RCH1pro::SnRK2.2 | *snrk2.2snrk2.3* | - | [43] | - | |
| *snrk2.2snrk2.3* SnRK2.2pro::SnRK2.2 | *snrk2.2snrk2.3* | - | [43] | - | |
| *aba2-1* | Col-0 | - | [64] | - | N156 |
| *cyp707a1cyp707a3* | Col-0 | - | [34] | - | |
| *cyp707a1cyp707a3* UBQ10pro::nlsABACUS2-400n – line 106 | *cyp707a1cyp707a3* | Floral dip | This study | FAST-RED | TBD |
| *cyp707a1cyp707a3* UBQ10pro::nlsABACUS2-400n - line 112 | *cyp707a1cyp707a3* | Floral dip | This study | FAST-RED | TBD |
| *cyp707a1cyp707a3* UBQ10pro::nlsABACUS2-100n line 5 | *cyp707a1cyp707a3* | Floral dip | This study | FAST-RED | TBD |
